# Passive thrust production in animal propulsion: exploitation of velocity gradients for energy conservation

**DOI:** 10.64898/2026.09.10.750785

**Authors:** Yangfan Zhang, Robin Thandiackal, Connor F. White, Jiacheng Guo, Yu Pan, Haibo Dong, George V. Lauder

## Abstract

Animals move through fluid environments that are rich in velocity gradients, yet how they exploit these flows to reduce locomotor costs remains elusive. We hypothesize that vertebrates utilize shear layers to passively produce thrust and establish a force equilibrium, thereby reducing locomotor cost. We discovered a shear-layer gait where fish preferentially position themselves within velocity gradients to maintain a steady posture and position at ∼50% of their sustained maximum speed. Compared to the matching speed of uniform flow, fish moved into an adjacent velocity gradient reduced active propulsion effort by 98% and metabolic costs by 74%, reaching a resting state. The mechanism is a force equilibrium between viscous drag on the high-velocity side of the body and anteriorly directed suction forces on the low-velocity side, fundamentally a passive thrust production driven by the pressure gradients. Such discovery has direct applications in bioinspired propulsion design and broad implications for movement ecology.

## INTRODUCTION

The ability to alter body movement patterns in response to different environmental conditions is fundamental to animal propulsion (*1*) (*2*). Without the capacity to modulate locomotion as conditions vary, animals would be confined to narrow ecological niches and constrained in locomotor performance. Animals continuously balance agility, speed and power to move through complex locomotor landscapes and to thrive in their ecological niches (*3*) (*4*) (*5*). These challenges have driven evolutionary diversification in morphology and physiology, yielding a wide variety of appendages, musculature, and cardiorespiratory systems that enable animals to adjust to environmental demands (*5*) (*6*) (*7*) (*8*) (*9*) (*10*) (*11*). These mechanical and physiological systems are intricately linked as cellular processes convert metabolic energy into mechanical work, which powers biomechanical structures to generate locomotor output and sustain movement.

Among the most prominent fluid dynamic features in locomotor landscapes is the presence of ubiquitous velocity gradients that arise in terrestrial, aerial, and aquatic habitats. The use of fluid velocity gradients by animals was first described by Leonardo da Vinci (*12*), yet after five centuries, our understanding of how animals modulate their kinematics to gain energetic benefits from ubiquitous velocity gradients remains incomplete. Velocity gradients are generated by the wake vortices of moving animals (*13*) (*14*), by flow around vegetation, rocks, and porous structures, and by boundary layers that form near solid surfaces (*15*) (*16*). Shear layers, which are regions of steep fluid velocity gradients, have attracted widespread attention in research on fluid mixing, nutrient transport, energy dissipation, habitat structure (*17*) (*18*) (*19*) (*20*) (*21*) (*22*), animal migration (*23*) (*24*), and animal abundance (*25*) (*26*). The ecological implications of fluid shear layers have been studied in hydraulic engineering, geophysics and fluid mechanics (*27*) (*28*) (*25*) (*29*) (*17*). Yet, whether and how animals can systematically exploit velocity gradients to reduce locomotor costs remains largely unresolved. In particular, it remains unknown whether vertebrates can use environmental shear layers not just to alter stability or maneuverability, but to generate thrust passively and achieve a net force balance that dramatically reduces, or nearly eliminates, the need for active propulsion.

Observations across taxa suggest that animals alter their locomotion in response to local fluid dynamic environments. Examples include the roller-coaster migration of bar-headed geese over the Himalayas (*30*), sharks riding updrafts produced by underwater channels and upslopes around atolls (*31*), fish altering kinematics when swimming in a drag wake (*32–34*), birds varying wing stroke patterns with changing air density (*35*) and during dynamic soaring (*36*, *37*), and seals gliding during dives with reduced locomotor movements and the resulting substantial energetic saving (*38*, *39*).

Although birds, fish, insects and crustaceans are all model organisms for understanding animal propulsion in a fluid, fish, especially those with undulatory, flexible bodies, are well-suited for studying fluid-mediated propulsion. The malleable, foil-shaped bodies of fish can interact with nearby vortices and velocity gradients (*40*) (*41*) (*42*) (*43*) (*44*) (*45*) (*26*) (*46*).

Importantly, aquatic locomotion is both energetically expensive and metabolically constrained because water is approximately 50 times more viscous and contains approximately 30 times less oxygen per unit of volume compared to air. Therefore, fish locomotion offers a unique model system to investigate fundamental principles of energy-saving during fluid-mediated propulsion and to study how animals might save locomotor energy by utilizing fluid velocity gradients.

We hypothesize that fish can position their bodies to passively generate thrust and balance drag forces resulting from the energy contained in fluid velocity gradients and thereby conserve energy compared to swimming in laminar flow conditions at the same free-stream velocity. Furthermore, we predict that a shear layer strength will exist where the net force on a flexible foil-shape approaches zero, and minimize the locomotor energy required for station holding.

To assess these hypotheses and predictions, we integrated behavioural characterizations, long-duration kinematic measurements, high-resolution respirometry, experimental and computational fluid dynamics (CFD), and force measurements on a fish model. We generated shear layers of various strengths in controlled conditions by positioning a foil at several angles in a recirculating flow tank, and compared the kinematics, energetics, and hydrodynamics of brook trout (*Salvelinus fontinalis*) that volitionally positioned themselves within shear layers to those of fish swimming in the free stream. This multifaceted approach allowed measurement of how fish select preferred locations in shear layers of various strengths and reduced their body and tail kinematics, how kinematics and bioenergetics are simultaneously coupled when exploiting a shear layer, and how the posture of fish reshaped the shear layer flow fields. We then used three-dimensional CFD based on trout kinematics and two-dimensional CFD simulations spanning a broad range of foil angles to reveal the flow physics that enable passive thrust generation.

These multifaceted analyses revealed that trout seek out and hold position within shear layers generated by the edges of angled foils, exhibiting a characteristic “shear-layer gait” in which body undulation is greatly dampened or nearly absent, even at flow speeds of ∼50% of their sustained maximum. In this gait, tail-beat amplitude and frequency are reduced by 81–91%, total kinematic effort falls by 45–98%, and the total energy expenditure is reduced by 71–74%, relative to free-stream locomotion at the same speed. Interestingly, fish position themselves *not* in the low-velocity drag wake but within the steep velocity gradient. At intermediate shear-layer strengths, a positive body angle of attack allows anteriorly directed force on the low-velocity side of the body to balance viscous drag on the high-velocity side, which produces a near-zero net force. Altogether, these results demonstrate that fish can exploit shear-layer velocity gradients as a passive thrust generator while achieving a force equilibrium that minimizes the energetic cost of locomotion to that of the overnight resting level. This mechanism links the physics of shear flows to animal movement ecology and provides a design principle for flow-sensing bio-inspired vehicles and robots that minimize the power consumption of movement by strategically positioning and orienting themselves within natural velocity gradients.

## RESULTS

### Fish positioning and locomotion kinematics in velocity gradients

Brook trout (*Salvelinus fontinalis*) consistently occupied positions lateral to angled foils (Fig.1), rather than locating throughout the flow field or in the drag wake behind the foil. When the foil was parallel to stream-wise flow (0°), fish locations were widely dispersed (95% area = 210 cm^2^; Fig. 1A, E). In contrast, with the foil at 45° and 90°, trout volitionally stationed lateral to both leading and trailing edges (95% area = 50 cm^2^), and rarely used the low-velocity drag wake immediately downstream of the foil. While holding position lateral to the 45° foil, fish exhibited a positive angle of attack of ∼10° relative to the incoming flow (Fig. 1B, D) and reduced positional variation compared to free-stream swimming (0° foil; Fig. 1H).

**Fig. 1.**
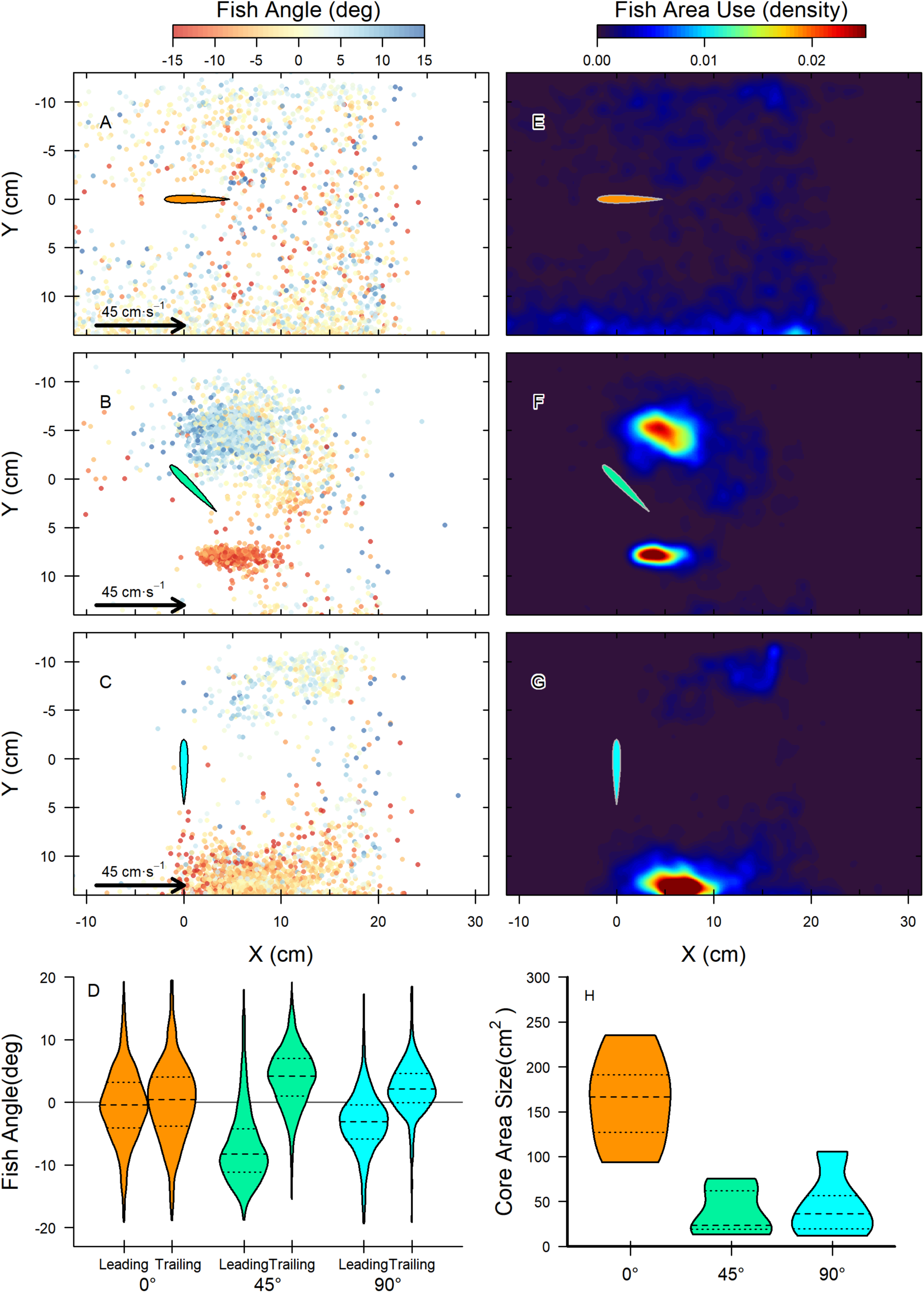
Fish location probability in flow fields. Volitional location and body angle (relative to the free stream) heatmaps for trout (*Salvelinus fontinalis*) swimming in the free stream with the foil at 0° (**A, E**), and in shear-layer velocity gradients generated by angling the foil to 45° (**B, F**) and 90° (**C, G**) relative to the streamwise flow. Fish preferentially position laterally to both the leading and trailing edges of the foil at 45 °, where shear layers are well-developed. With the foil at 90° and generating a wide wake, fish locate primarily to one side of the foil and near the flow tank wall, but this behaviour did not occur when a shear layer was generated by an angled foil at 45°(**F**). Probability distributions in **E, F, G** were calculated using Gaussian kernel density estimation. Fish body angle varied with foil angle (**D**), and body location was more restricted when fish held station to the leading and trailing edge of the foil. The length of the foil is 6.8 cm.

Trout located within velocity gradients lateral to 30° and 45° foils (suppl. movies 1 & 2) exhibited greatly dampened to nearly absent body undulation with sharply reduced tail-beat frequency and amplitude, while maintaining a positive body angle of attack, and held station in oncoming flow (at 50% of sustained maximum speed). We term this locomotor pattern a “shear-layer gait” (Fig. 2C-E), which contrasts with the classic undulatory deformation of the body during free-stream swimming (0° foil; Fig. 2F). When trout occupied the shear layer generated by 45° and 30° foils, average tail beat frequency (*f*_tail_) was 1.9 Hz or 0.45 Hz, and an average tail beat amplitude (Amp_tail_) was 0.06 BL or 0.035 body length (BL) respectively, which were 64–91 % lower than *f*_tail_ (average 5.4 Hz; *F*_3,31_ *=* 157.3, *p <* 0.001), and 63–81 % lower than Amp_tail_ (average 0.16 BL; *F*_3,31_ *=* 201.0, *p <* 0.001; Fig.2 C-F, H, I) in the free stream. Amp_tail_ of fish swimming in the wake of 90° foil was statistically indistinguishable from that of freestream swimming (average = 0.15 BL; *F*_3,31_ = 201.01, *p* = 0.978), but the *f*_tail_ was 55% lower (*F*_3,31_ *=* 157.3, *p* < 0.001). Overall, total kinematic effort of the shear-layer gait, estimated as Amp_tail_ • *f*_tail_, was 45–98% lower than free-stream swimming (0.82 BL·s^-1^) at 50% of sustained maximum speed (Fig. 2H: 90° = 0.36 BL·sec^-1^, 45° = 0.12 BL·sec^-1^,30° = 0.02 BL·sec^-1^; *F*_3,31_ *=* 394.8, *p <* 0.001; suppl. movie 2).

**Fig. 2.**
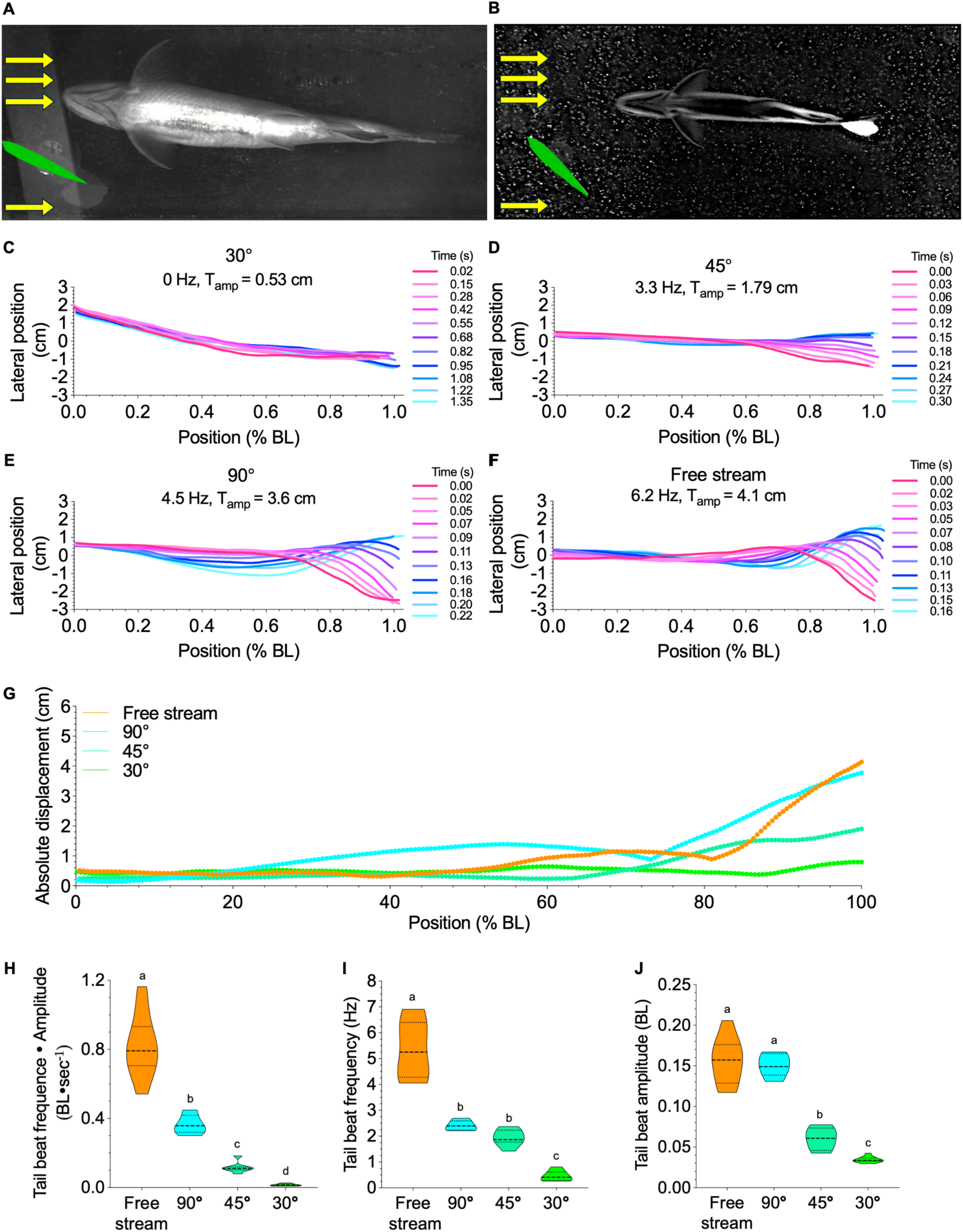
Kinematics of trout (*Salvelinus fontinalis*) swimming in shear layers of different strengths resulting from altering foil angle. (A) Images from light video and (B) particle image velocimetry recordings of fish swimming within a shear layer created by a foil (green shape) angled at 30° and 45° to the streamwise flow (yellow arrows). (C-F) Midline kinematics of fish interacting with different strengths of velocity gradients produced by foils angled at 30°, 45°, 90° and 0° (free-stream flow with the foil present) to the streamwise flow. (G) Amplitude of lateral midline motion for fish in shear layers at each foil angle. (H-J) Statistical comparisons of kinematic effort (tail beat frequency × amplitude), tail beat frequency and amplitude when fish interact with velocity gradients generated by foils at 30°, 45°, 90° and 0° to the streamwise flow. Foil chord length is 6.8 cm.

Kinematic effort of the shear-layer gait differed among foil angles. Fish shear-layer gaiting near the 30° & 45° foils had a 68–96% lower total kinematic effort than that near the 90° foil (0.36 BL·s^-1^; *F*_3,31_ *=* 394.8, *p <* 0.001), driven by 60–77% lower Amp_tail_ (90° = 0.15 BL, 45° = 0.06,30° = 0.035,; *F*_3,31_ *=* 201.0, *p <* 0.001). Although fish shear-layer gaiting near the 45° foil had the same *f*_tail_ as that of the 90° foil, fish shear-layer gaiting by a 30° foil had 81% lower *f*_tail_ (30° = 0.45Hz, 90° = 2.4 Hz; *F*_3,31_ *=* 157.3, *p <* 0.001; Fig. 2E) than trout lateral to the 90° foil.

### Experimental fluid dynamics of the shear-layer gait

Flow around each side of the foil edges generated steep velocity gradients. At the leading edge of the 45° and 90° foil, free-stream velocity reached 0.57 m s^-1^, but dropped 90% to a minimum streamwise velocity of 0.055 m s^-1^ in the drag wake behind the foil (suppl. movie 3a, b, c). The sharp velocity gradient from the free-stream to wake region formed a streamwise shear layer as the oncoming flow moved downstream of the foil (Fig. 3; suppl. movies 3a, 3b, 3c).

**Fig. 3.**
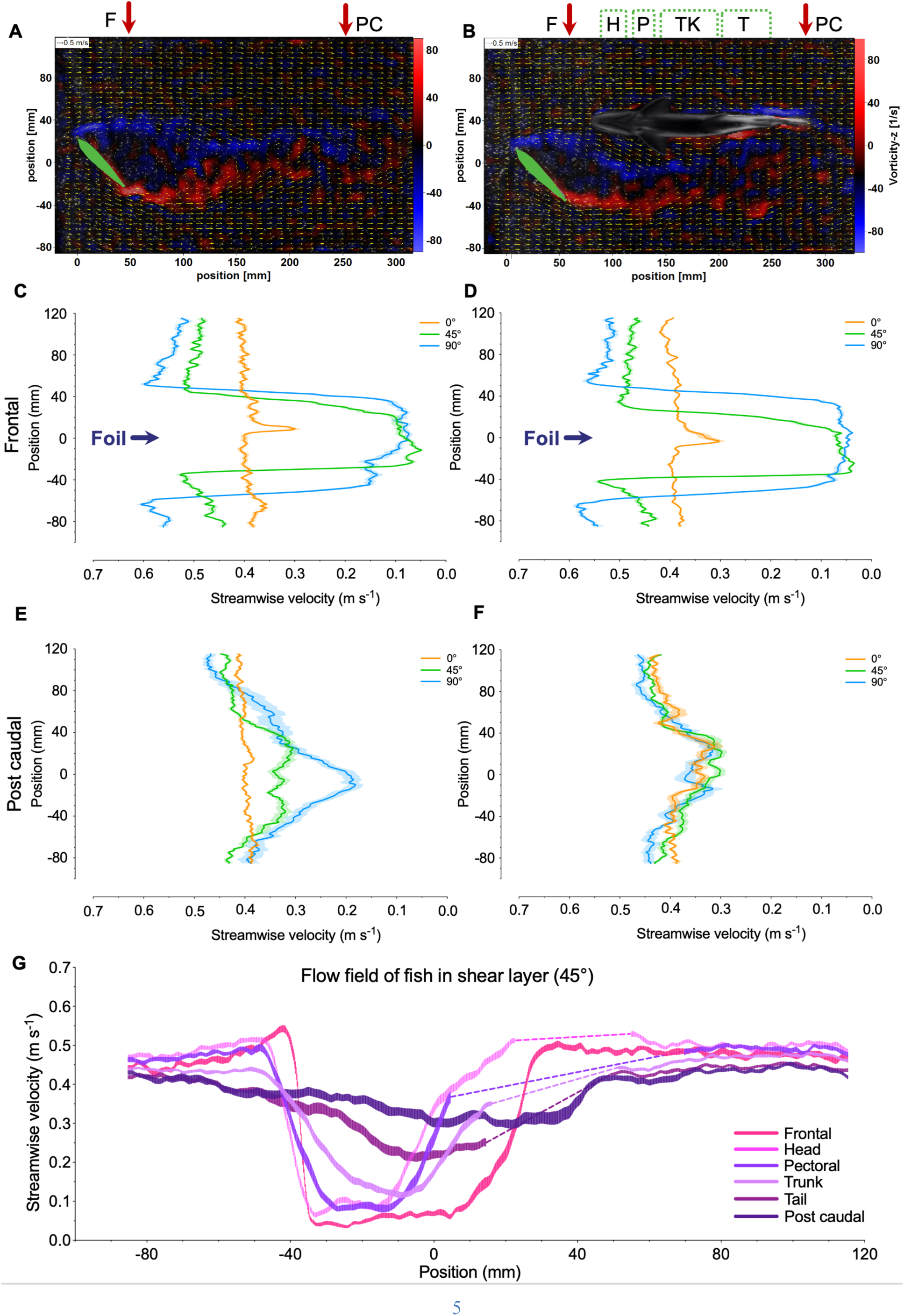
Particle image velocimetry (PIV) analyses of control and shear layer locomotion. Flow fields generated by a foil angled at 0°, 45°, and 90° to the streamwise flow are shown for control conditions without a fish (**A**) and with a fish **(B**) holding position in the shear layer laterally behind the leading edge of the foil. Background colour indicates vorticity, and yellow vectors indicate flow velocity. Velocity profiles were measured transverse to the free stream at frontal (F), head (H), pectoral (P), trunk (TK), tail (T) and post-caudal (PC) locations, as indicated in (**B**). Panels **C** and **E** show the flow field under control conditions, while panels (**D)** and (**F)** show profiles with a fish present in the shear layer. (**G**) Cross-sectional profiles of streamwise flow at the indicated locations (dashed lines indicate the location of the fish body). Shaded regions represent the standard error of the mean. Foil chord length is 6.8 cm.

The presence of the fish did not substantially alter the flow field directly behind the foil, but it dampened the steep drop in the velocity profile over streamwise direction: the velocity gradient became shallower as vortices rolled along the side of the trout body and reached the 28% velocity reduction (from free stream) at post-caudal region (Fig. 3F), compared to that of 90% velocity reduction in the drag wake (Fig. 3D). The velocity gradient profile in the post-caudal region of the fish was similar regardless of foil angle (Fig. 3F). However, at the same post-caudal location in the flow field but without fish, the velocity at the 90° foil dropped by 52% relative to that of 0° foil (Fig. 3E). The fluid field demonstrated that trout positioned themselves within streamwise shear layer (Fig. 3G, dashed lines indicate fish position; suppl. movie 3a&b).

Shear-layer strength, quantified as the velocity gradient across a distance (m·s⁻¹·mm⁻¹), depended on foil angle and streamwise distance (Fig. 4). At a vertical profile aligned at the center of the foil (Fig. 4: blue vertical line), shear-layer strength increased with foil angle (30°: 0.044 ± 0.0016 m·s^-1^·mm^-1^, 45°: 0.037 ± 0.0031 m·s^-1^·mm^-1^, 90°: 0.074 ± 0.0053 m·s^-1^·mm^-1^; *F*_3,68_ *=* 132.6, *p <* 0.001; Fig. 4), where shear-layer strength was similar between the 30° and 45° foil (*F*_3,68_ *=* 132.6, *p =* 0.293). However, the variability of shear-layer strength at the centre of 30° foil (C.V. = 10.7%) was lower than that of the 90° and 45° foils (C.V.: 90° = 22.7%, 45°= 26.5%). Moreover, shear-layer strength declined over streamwise distance for all angles, reaching 0.0081, 0.020 and 0.0070 m·s^-1^·mm^-1^ at 120 mm downstream for 30°, 45° and 90° foils, respectively, which were all significantly lower than shear-layer strength near the foil (*F*_1,68_ *=* 279.3, *p <* 0.001). At 120 mm downstream, the 45° foil produced a 143-183% higher shear-layer strength than that of the other angles (*F*_3,68_ *=* 10.9, *p ≤* 0.011; Fig. 4B).

**Fig. 4.**
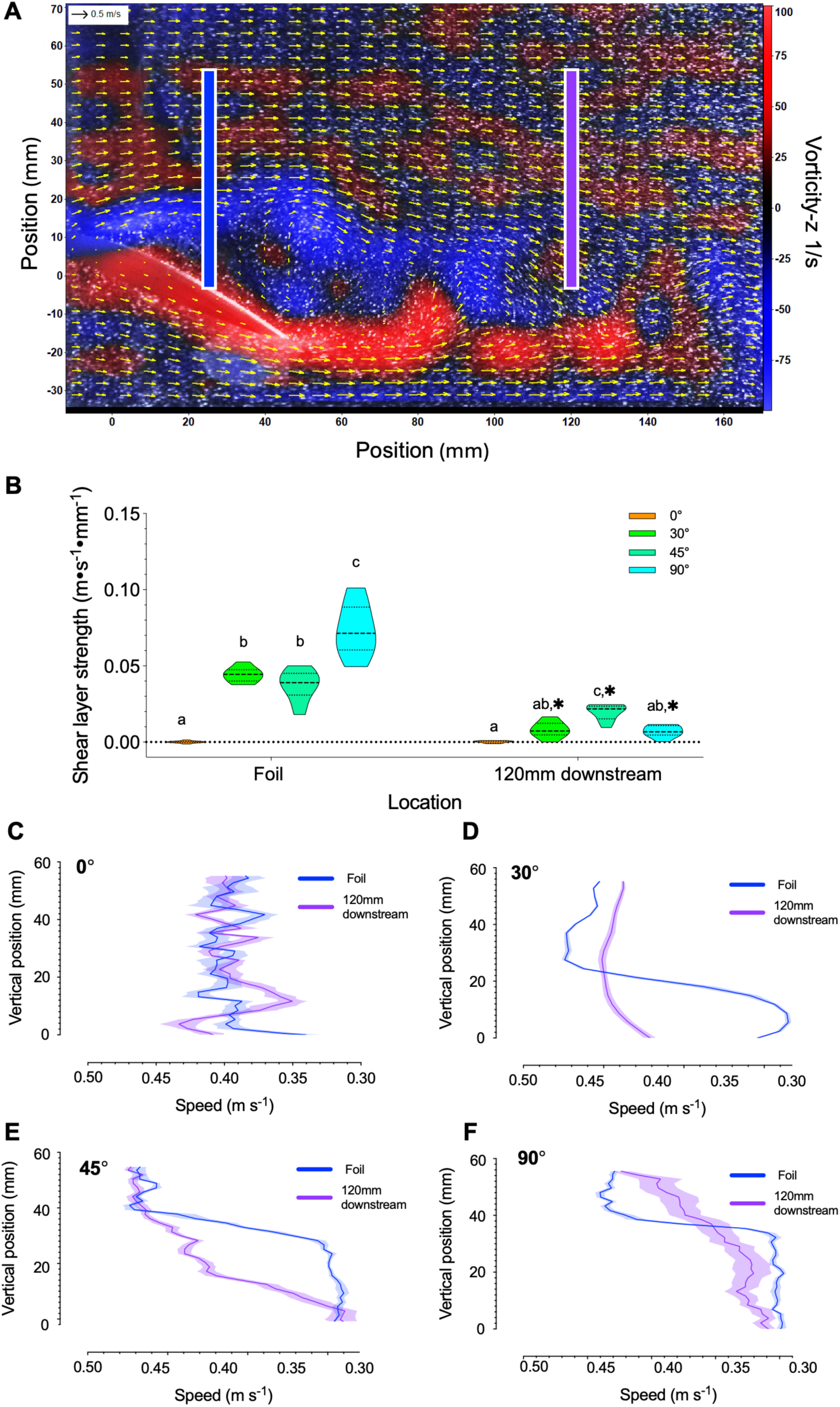
Particle image velocimetry (PIV) analyses of shear-layer strength. Shear-layer strength is controlled by orienting a foil at 0°, 30°, 45°, and 90° to the streamwise flow, and measured under control conditions without fish (**A**). Blue and purple vertical lines denote the cross-sectional profile locations (55 mm in length) sampled at the centre of the foil and ∼120 mm downstream. Shear-layer strength is defined as the velocity gradient (the rate of change in velocity with distance) across the cross section. **(B**) Violin plots of shear-layer strength across four foil angles and two locations. (**C–F**) Velocity profiles of the shear layer region and a section of the freestream (40–55 mm position) are shown as a reference across four foil angles and two locations. Shaded regions indicate the standard error of the mean; Shear layers generated by the 90° foil exhibit greater temporal variation in strength. Letters denote differences in shear-layer strength among foil angles at the same location. Asterisks indicate significant differences in shear-layer strength between locations for a given foil angle. Foil chord length is 6.8 cm.

### Metabolic cost of shear layer locomotion

Direct measurements of whole-animal metabolic rate showed that trout exhibiting a shear-layer gait achieved large energy saving (71–74% lower) compared to free-stream locomotion (Fig. 5). Aerobic metabolic rate of the shear-layer gait was statistically indistinguishable from the overnight resting value (shear-layer gait vs rest: 73 vs 84 mg O_2_ h^-1^ kg^-1^ or 3.96 vs 4.55 kJ kg^-1^ min^-1^; *p* ≥ 0.142, Fig. 5A), which was 71% lower than the aerobic costs of locomotion (254.9 mg O_2_ h^-1^ kg^-1^; *F*_1.072, 3.216_ = 539.2, *p* = 0.006) in the same free-stream velocity. Total locomotor energy expenditure (aerobic metabolism plus excess post-exercise oxygen consumption, EPOC) was reduced by 74% in the shear-layer gait relative to free-stream swimming (15.3 kJ kg^-1^ min^-1^; *F*_1.111, 3.334_ = 303.4, *p* = 0.001) (Fig. 5 A−D).

**Fig. 5.**
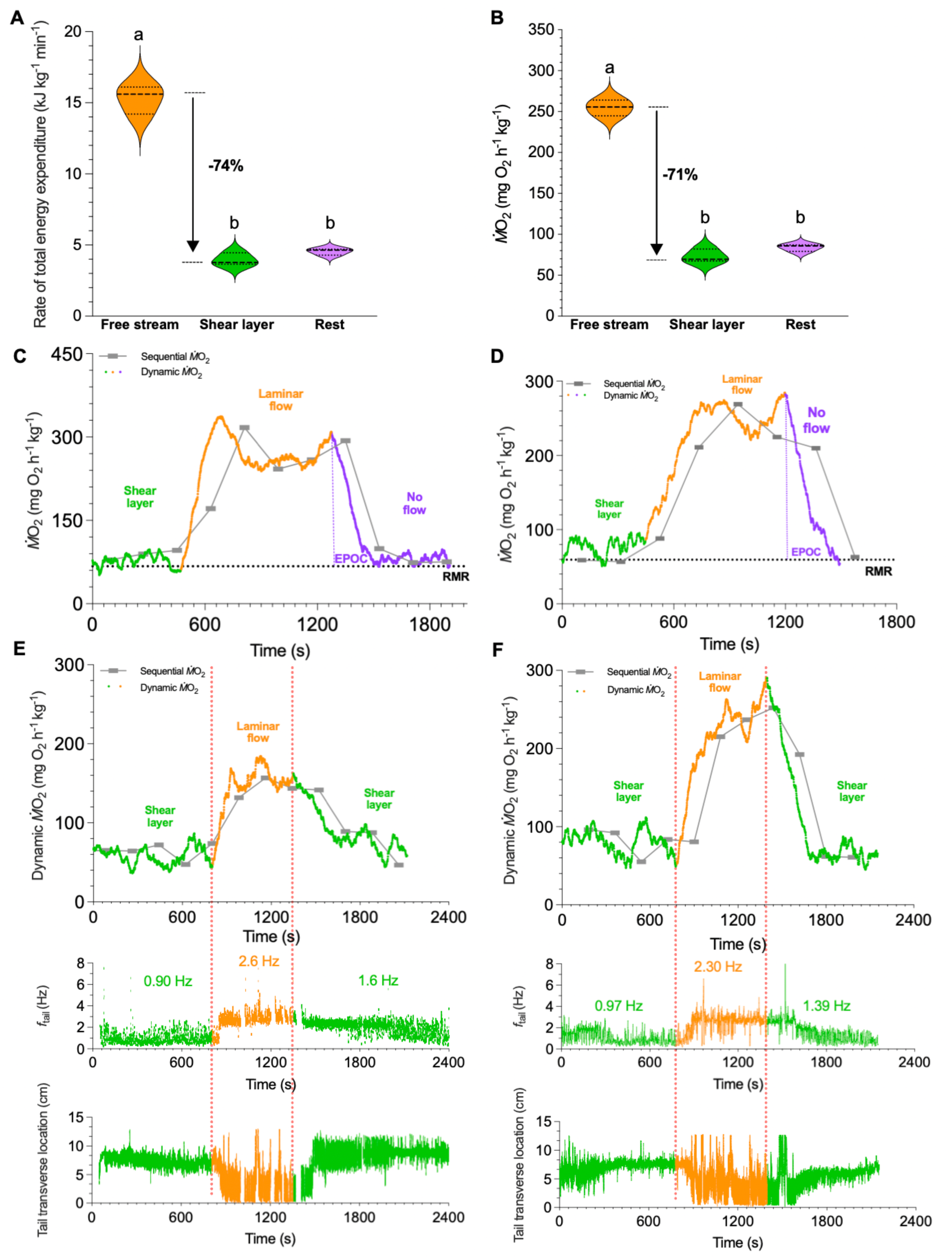
Steady-state and dynamic metabolic rate during locomotion in a shear layer. Total energy expenditure, including both aerobic and non-aerobic (**A)** and aerobic metabolic rate **(B**) of fish locomotion in free stream (0°), in a shear layer (30° foil angle), and at rest. (**C, D**) Time series of metabolic rate (*Ṁ*O_2_) during swimming in a shear layer, in free stream, and in no-flow conditions, as the foil generating the shear layer was rotated from 30° to 0°. The no-flow condition enables measurement of non-aerobic costs of locomotion using excess post-exercise oxygen consumption (EPOC). Combined aerobic swimming costs and EPOC provide an estimate of the total energetic cost. Conventional analyses of sequential metabolic rate measurements are provided as a reference (grey curve and bars; *see* Methods). (**E, F**) Synchronous measurements of metabolic rate and kinematic modulation (tail beat frequency and transverse location) as fish transition between shear-layer and free-stream flow fields, as denoted by the dotted lines. Fish substantially reduce both the tail transverse location and beat frequency (the average values are labelled) of body movements when holding position in the shear layer.

The dynamic coupling between kinematics and metabolism was studied by rotating the foil within the respirometer to alternately generate and remove the shear layer while continuously recording aerobic metabolic rate and tail kinematics. The dynamic aerobic metabolic rate of shear-layer gait oscillated around 68 mg O_2_ kg^-1^ h^-1^ accompanied by *f*_tail_ ranging from 0.38 to 2.1 Hz. When the foil was rotated from 30° to 0°, eliminating the shear layer, aerobic metabolic rate rose steeply by 193% to ∼200 mg O_2_ kg^-1^ h^-1^ and *f*_tail_ increased to 2.4–3.1 Hz as fish transitioned to undulatory swimming in free stream. When the foil was rotated back to 30°, restoring the shear layer, fish re-entered velocity gradients and re-established the shear-layer gait, where *f*_tail_ decreased to 0.63–2.33 Hz, and the aerobic metabolic rate was reduced by 68% to ∼64 mg O_2_ kg^-1^ h^-1^ (Fig. 5E, F). Tail kinematics during the shear-layer gait were also less variable than in the free stream (coefficient of variation of *f*_tail_: 21% vs 59%). These synchronized measurements demonstrated a tight temporal linkage between shear layer flow dynamics, kinematic modulation and locomotor energetic costs.

### Force measurement on a trout body model

To understand the spatial distribution of hydrodynamic forces, we measured streamwise (X), lateral (Y), and vertical (Z) forces and torques on a 3D-printed trout model (across 112 grid points) downstream of 45° and 90° foils (Fig. 6 A,E,I,M). When located downstream and inline with the angled foil, the model experienced net thrust (maximum *F*_x_ = 155 mN, Fig. 6) in the area that extended 15.5 cm downstream and 5 cm laterally to the 90° foil (Fig. 6A), which exceeded the area and thrust magnitude behind the 45° foil (Fig. 6E, downstream = 12.5cm, laterally = 3cm, maximum *F*_x_ = 106 mN). Lateral forces (*F*_y_) were minimized inline with the 90° foil (Fig 6B), whereas the centre-line lateral equilibrium of a 45° foil emerged behind the trailing edge (Fig. 6 F). When the fish model was at 0° angle of attack, the *F*_x_ forces were minimized towards the lateral edges of the foil, and *F*_y_ forces were minimized behind the foil, which created a narrow area of net-zero forces (Fig. 6 D,H).

**Fig. 6.**
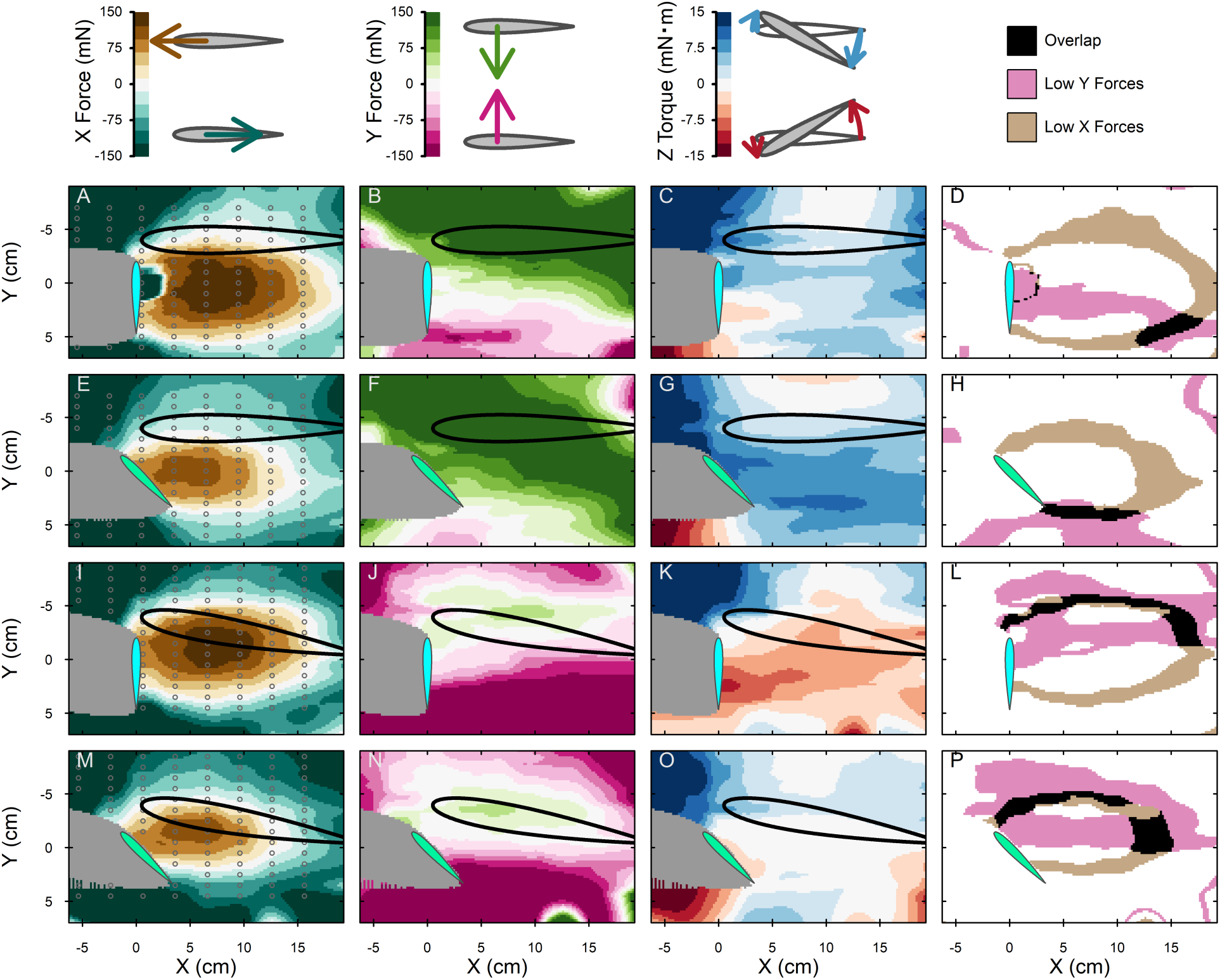
Spatial variation in forces acting on a fish model behind a 45° and 90° foil. Spatial distributions of streamwise force (X-force; **A, E, I, M**), lateral force (Y-force; **B, F, J, N**), and yaw torque (Z-torque; **C, G, K, O**) on a fish model (black outline) located downstream of foils angled at 45° and 90°. Icons above each column indicate the direction of the forces and torques. Each dot indicates a location at which forces and torques were measured behind the foil (cord length = 6.8 cm). Forces were measured surrounding a 90° (**A-D** and **I-L**) and 45° foil (**E-H** and **M-P**). The fish body was oriented at 0° (**A-H**) and 10° (**I-J**) to the flow. Areas of low net X & Y forces (<30 mN, **D,H,L,P**) highlight the regions (black area) where the model experiences low net forces.

When the model fish orientation matched that of live fish (∼10° angle of attack, see Fig 1), a region of net-zero force occurred in a location that resembled the preferred position of biological fish (Fig. 6. D,H). Although the maximum thrust changed <15%, the area of net upstream forces was shifted 2 cm laterally (Fig.6 D to L & H to P), and the area of low lateral forces was substantially shifted (Fig. 6 J,N). This generated a substantially expanded region of low net forces (266% for 45° foil, 223% for 90° foil; Fig.6 L,P) that overlapped with areas of low yaw torque (Fig. 6 K, O).

The stability of thrust (*F*_x_) regions at the leading and trailing edges varied, despite both experiencing low net forces. Any deviation in position of the trailing edge accumulated the force that destabilized the fish, whereas the thrust region at the leading edge stabilized the fish, as forces returned the fish towards their original low-force position. Generally, the medial and leading edge of the low lateral (*F*_y_) region were stabilizing. Considering all forces on the model fish together, a stable fluid corridor is evident at the leading-lateral edge, which corresponded to the *in vivo* position and posture of the fish.

### Computational fluid dynamics of the shear-layer gait

Three-dimensional computational fluid dynamics (CFD) simulations reconstructed from videography of trout exhibiting the shear-layer gait lateral to a foil clarified the physical mechanisms underlying the force balance on the fish body (Fig. 7; suppl. movies 4a,b).

**Fig. 7.**
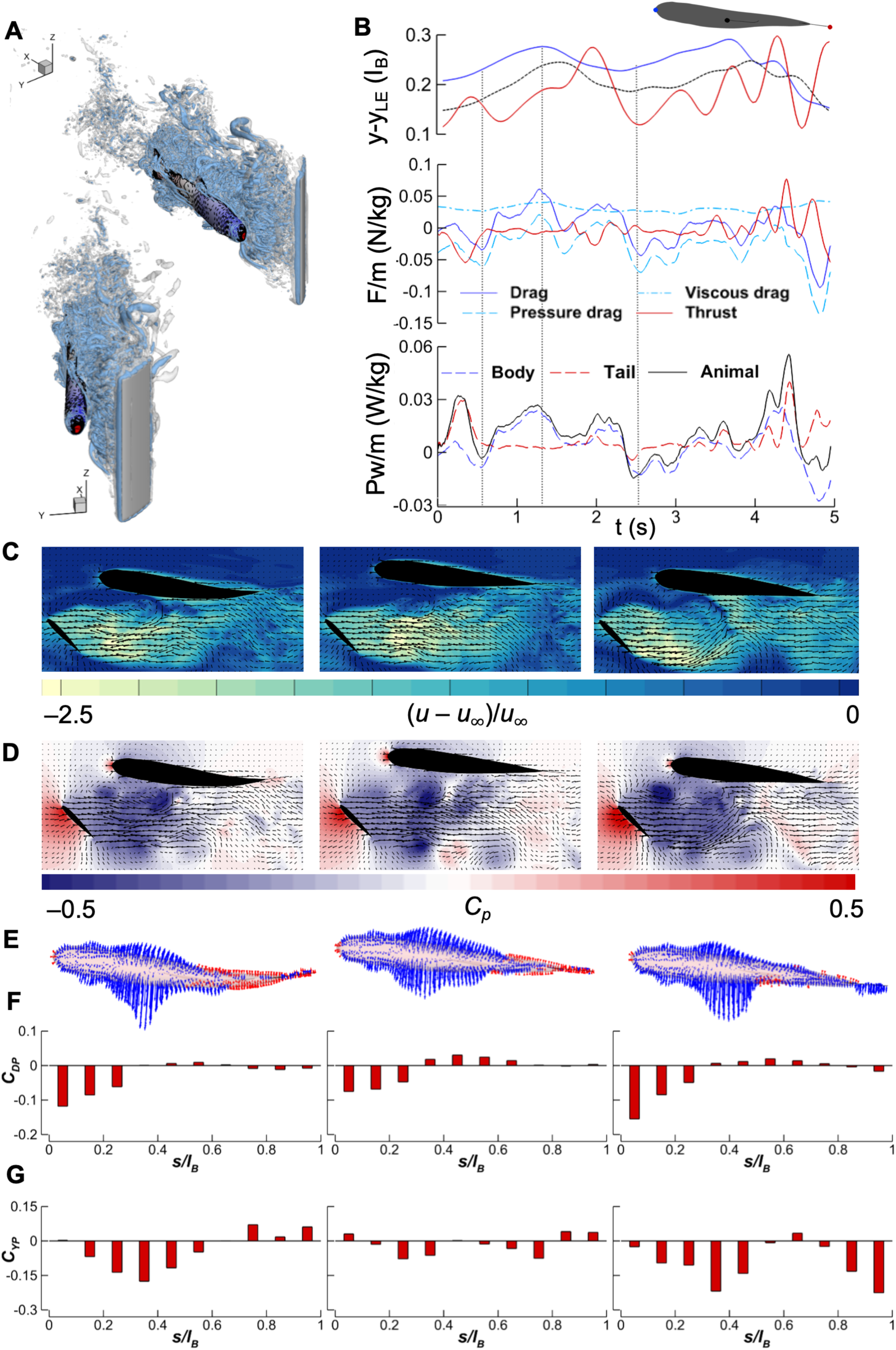
Three-dimensional computational fluid dynamics (CFD) of steady station holding in the wake of the 45° foil. (**A**) Three-dimensional wake structure visualized using Q-criterion isosurfaces in isometric and frontal views. (**B**) The instantaneous lateral positions, streamwise forces, and hydrodynamic power consumption during the simulation window for station holding. To reveal the underlying flow mechanics, three representative instances demonstrate the profiles when body drag is low, high, and low (**C, D, E, F, G**). Flow dynamics and body forces at the three instants: (**C**) streamwise flow profile with flow velocity vectors (*u*_∞_ is the background flow speed), (**D**) distribution of pressure (*C_P_*) in the flow, (**E**) pressure force vectors acting on the fish (blue: suction; red: positive pressure), (**F**) distribution of profile drag coefficient (*C_DP_*), and (**G**) distribution of pressure lift force coefficient (*C*_F*P*_) (*C*_F*P*_ < 0 representing lateral force directed toward the centre of the foil wake).

Simulations revealed that fish experienced persistent viscous drag (Fig. 7B), but pressure drag coefficient was periodically zero or negative and fluctuated more strongly than the viscous drag, indicating an anterior suction force (Fig. 7B). The total drag coefficient, including both viscous and pressure drag, oscillated around zero, which enabled an overall force balance for fish in the shear layer (Fig. 7B). Hydrodynamic power consumption tracked fluctuations in total drag and remained low throughout the simulation.

Three representative instances illustrate flow interactions that enabled drag and power consumption reductions by the shear-layer gait (Fig. 7C, D). The trout body was oriented at an angle of attack (∼10°) in the velocity gradient lateral to the foil edge and well outside the suction region behind the center of the foil (Fig. 7D). The side of the body facing the drag wake behind the foil experienced reduced velocity (Fig. 7C) and pressure (Fig. 7D), which generated an anteriorly-orientated force on the first ∼35% the body and minimized yaw torque (*34*)), while the opposite side experienced posteriorly-orientated forces (Fig. 7E,F). This suction force persisted even at a moment of high drag (Fig. 7F mid-panel). As fish momentarily approached the drag wake, the suction force also applied a stronger lateral component (Fig. 7G left & right-panels vs 7G mid-panel). The net effect balanced forward suction and backward viscous drag (Fig. 7F, G) with a minimum yaw torque, which yielded a near-zero time-averaged drag and low hydrodynamic power.

To examine the effects of shear-layer geometry on the force balance, we performed two-dimensional CFD simulations spanning foil angles from 0° to 90° and a fish model at 10° (Fig. 8A, suppl. movie 5). The drag force decreased with increasing foil angle (*F*_6,13993_ = 6456, *p* < 0.0001), and switched from downstream (drag, + C_d_) to upstream (thrust, - C_d_) orientation between 30° and 45° foil (Fig 8C). Lift exhibited similar trends as the foil angle increased (*F*_6,13993_ = 6044, *p* < 0.0001; Fig.8D). The lateral forces were inversely related to foil angle. The net lateral forces reversed from pointing away from the foil to pointing towards the foil between 45°–60°. Temporal variance in the streamwise and lateral forces rose with foil angle (Fig 8B).

**Fig. 8.**
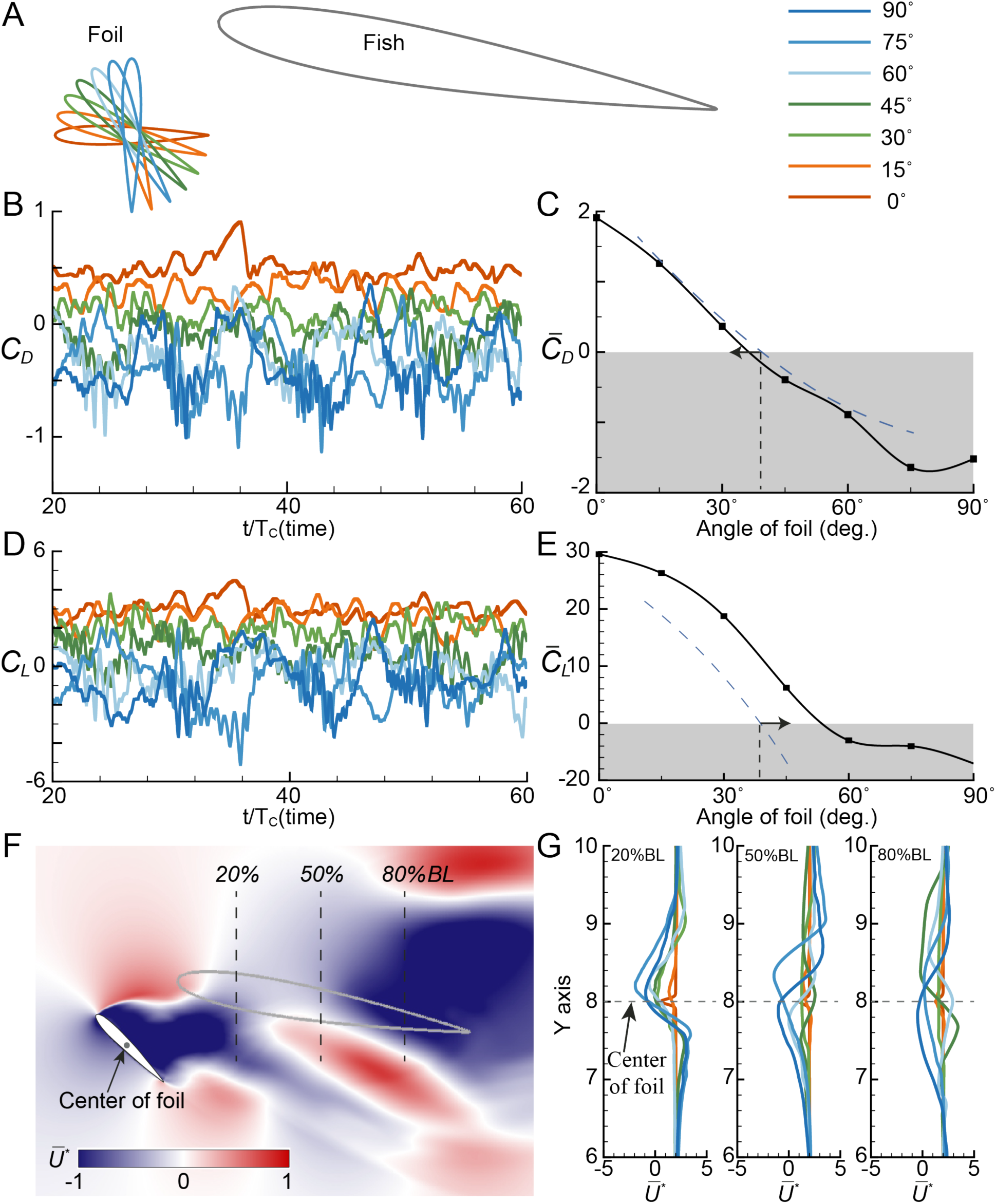
Two-dimensional computational fluid dynamics simulation of various foil angles and their interaction with a fish body model. (**A**) Schematic of the relative position of the foil with respect to the model fish. The centers of the foil and fish body remain fixed, while the foil angle rotates from 0° to 90° in 15° increments. The foil angle is defined as the angle between the foil chord and the oncoming flow. (B) Time history of the drag coefficient CD of the fish as a function of foil angles. (C) Time-averaged drag coefficient ‘C’D’ of the fish body as a function of foil angle. The time-averaged drag on the fish body decreases as the foil angle increases, but approaches zero at ∼40°, consistent with experimental observations. Because the simulations are two-dimensional and at a lower Reynolds number (Re = 5000), the computed drag is lower than the actual drag (dashed line). (D) Time series of the lift coefficient CD of fish at different foil angles. (E) Time-averaged lift ‘C’L’ on the fish body decreases as the foil angle increases. At ∼40°, ‘C’L’ approaches zero, implying that the fish can maintain lateral stability with minimal energy expenditure. Due to the model simplification and lower Reynolds number, the computed ‘C’L’ is higher than the actual value, but the qualitative differences remain consistent. (F) Mean flow behind a 40° foil. A virtual fish (grey foil outline) is placed downstream of the foil wake to examine the effects of the foil on the local fluid field around the fish at a normalized velocity (U*∗). (G) Time-averaged velocity profiles U*∗ at 20%, 50%, and 80% body length (from the leading edge of the fish for foils at different angles). The oscillations in flow are consistent with small oscillatory movements of the fish body when stationed in the shear layer.

Force fluctuation, quantified by the root-mean-square of drag (RMS) (C_D_) and lift (C_L_), and their time derivatives, also increased with foil angle (Fig. S2). The RMS of drag force amplitude was ∼0.10 at 0°–30° foil, increased to ∼0.16 at 45°, and reached ∼0.27 at 90° (Fig.8B). The RMS of the time derivative of drag increased from 0.42 at 0°, to 0.67 at 30°, 1.22 at 45° and 1.64 at 90°.

These results suggested that maximum energetic benefit is achieved at the 30°–40° foil angles, where the time-averaged drag coefficient was close to zero, and force fluctuations were modest (Fig. 8C). Under these conditions, the simulated fish experienced reduced to near-zero net forces, consistent with the low kinematic effort, near-resting metabolic costs, and net-force equilibrium measured experimentally for shear-layer gait exhibited at 30°–40° foils.

## DISCUSSION

Animals use fluid shear layers for sensing, orientation, and movement. For example, plankton luminescence responds to the intensity of fluid shear over swimming marine mammals (*47*). Copepods orient and track odour plumes driven by oceanic shear layers (*23*). Modelling approaches suggest that oceanic birds achieve energy-neutral wind–shear soaring using velocity gradients formed by air movement over ocean waves (*48*) (*49*). Although substantial reductions in wing beat frequency and heart rate support the hypothesis of energy saving in birds (*50*), the direct metabolic verification of energy saving by birds in atmospheric shear layers is understandably challenging. Thus, the kinematic and fluid dynamic mechanisms by which animals utilize fluid velocity gradients to save energy remain a largely unexplored area of organismal propulsion with broad implications for ecological, evolutionary and engineering aspects of movement.

Here we reveal the mechanisms involved in locomotor energy saving when fish exploit near-obstacle shear layers. Fish position and orient their foil-shaped body within the shear layer produced by an obstacle at a positive angle of attack to passively generate thrust and achieve a net force equilibrium, allowing station holding in flows of 50% of their sustained maximum speed while decreasing total kinematic effort by up to 98% and reducing metabolic cost by 74%, reached that of overnight resting levels (Fig. 5A). This level of metabolic energy saving is greater than when fish utilize vortex shedding with the Kármán gait at a similar free-stream flow velocity (a 53% reduction in locomotor cost) (*51*).

Across integrated behavioural, kinematic, energetic, force measurements, and experimental and computational analyses of fluid dynamics (Fig. 9A, B), we describe a distinct locomotion strategy: a “shear-layer gait” (suppl. movies 1–3). This locomotor pattern exhibits the following essential features: 1) animals volitionally occupy steep velocity gradients; 2) a small angle of attack of the body generates lift; 3) body movement and energetic costs are substantially reduced; and 4) the body experiences near-zero net fluid dynamic forces and torques.

**Fig. 9.**
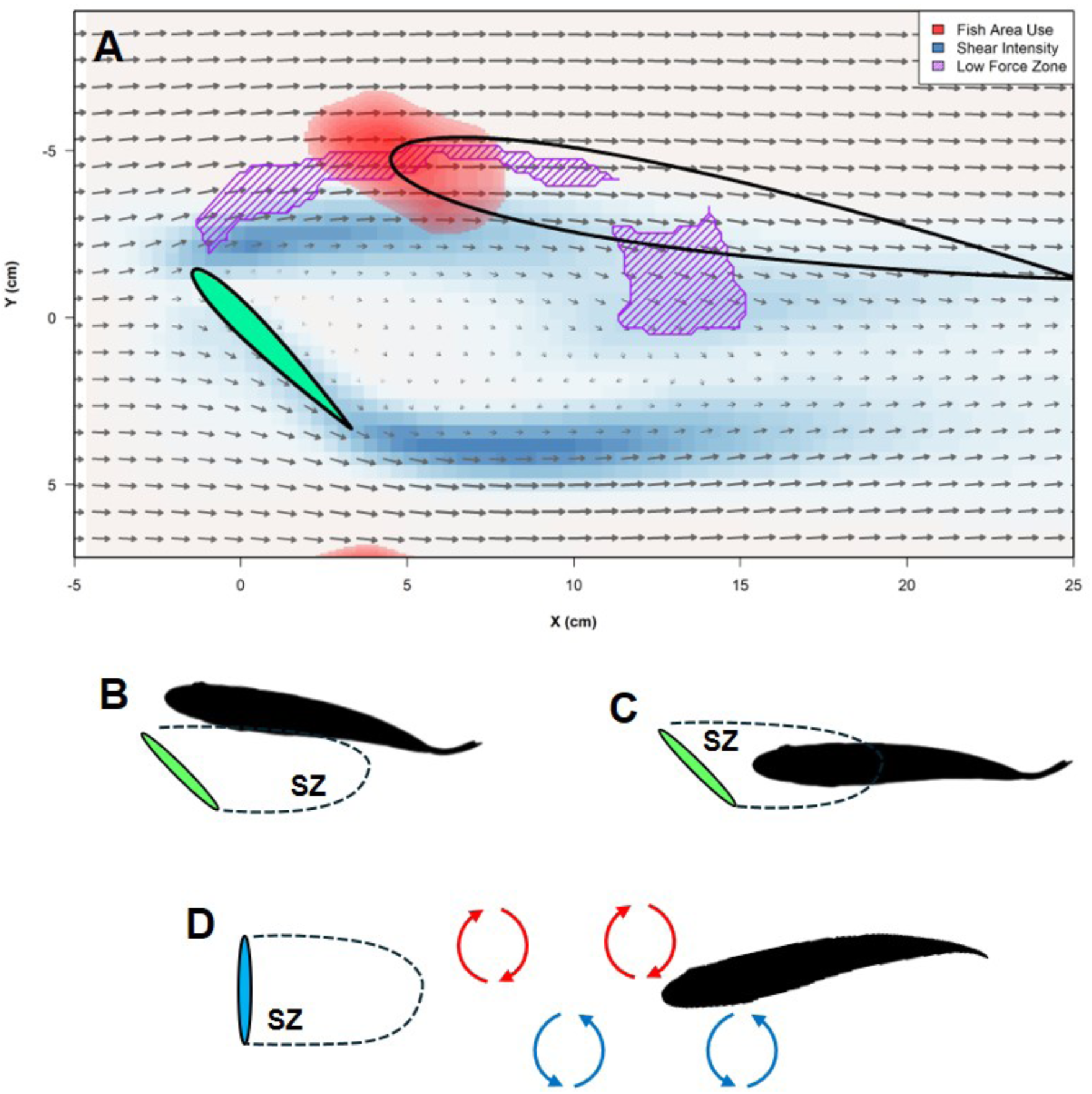
Summary of the shear-layer gait and comparison with other mechanisms of locomotor energy conservation. (**A**) The shear-layer gait is used when fish volitionally position themselves within a region of a steep velocity gradient formed between freestream flow and the wake of an obstacle, and achieve an overall force balance on the body, allowing station holding with greatly reduced energetic cost compared to free-stream locomotion. Schematic diagrams to compare and contrast fish positioning within a shear layer generated by a foil oriented at 45° to free stream flow (**B**), fish locating within the suction zone (SZ) in the region downstream near the foil, shown here at a 45° angle (**C**), and fish exhibiting the Kármán gait (**D**) well downstream of the suction zone where passive body deformation allows station keeping by interacting with the Kármán vortex street (alternating vortices denoted by red and blue arrows) shed from the foil edges. A 90° foil angle is shown as most Kármán locomotion studies have used either a D-cylinder or a round cylinder to generate the vortex wake. Fish positioning in all three locations can save substantial amounts of energy compared to locomotion in the free stream.

Mechanistically, the shear-layer gait arises from a precise balance between viscous drag on the high-velocity side of the body and anteriorly directed suction forces on the low-velocity side, which is fundamentally driven by the velocity and pressure gradients within the shear layer. The experimental and computational results illuminate a previously underappreciated mechanism by which a foil-shaped body can dramatically reduce the energetic cost of experiencing high-energy fluid environments and offer a unifying principle for efficiently moving through fluids: appropriate kinematics can harness external fluid kinetic energy to passively generate thrust.

### Mechanisms of locomotor energy conservation

Aquatic animals can conserve locomotor energy in rapid flow environments through at least three broadly recognized strategies: (1) entrainment in near-obstacle flow patterns (*46*) (*26*, *52*), (2) energy extraction from a Kármán vortex street downstream of an obstacle (*34*, *53*) (*33*), and (3) use of the boundary layer near solid surfaces (*54*, *55*). All three rely on exploitation of spatiotemporal variation in flow velocity (Fig. 9).

Entrainment involves station holding near objects in flowing water, as coined by Sutterlin and Waddy in 1975 (*56*), who suggested that entrainment could possibly allow “fish to maintain position with minimum expenditure of energy.” Fish can entrain at a suction zone behind an obstacle extending approximately two obstacle diameters downstream, where flow velocity is reduced. Fish locating with their head within the suction zone experience an upstream force that is countered by drag on the region of the body located outside the suction zone. Fish can also entrain in the bow wake, in front of obstacles, and benefit from the pressure gradient formed by flow impacting the obstacle (*51*) (*26*). Previous research on fish entraining at the edges of obstacles showed a reduced tail beat frequency (*f*_tail_) that is similar to that occurring in the experiments reported here with a 90° foil with a shear layer strength of 0.074 m·s^-1^·mm^-1^ (*56*) (*52*) (*26*) (*51*). Obstacles of different configurations can generate shear-layer zones of different strength and stability that alter fish locomotor energetic costs and dynamics (*e.g.,* midline kinematics, fin movement).

Kármán gait locomotion, in contrast, occurs in the alternating Kármán vortex street downstream of the suction zone behind an obstacle (*33*) (*34*) (Fig. 9D). Trout swimming in a Kármán vortex street locate well downstream of the suction zone and increase tail beat amplitude (Amp_tail_) and match *f*_tail_ to the vortex shedding frequency behind the obstacle. Body movement is largely passive with reduced muscular activity, which reduces locomotor energy expenditure up to 53% compared to swimming in the free-stream (*34*).

Fish can also benefit from greatly reduced drag when locating either partially or completely within boundary layers where flow varies from zero at the surface due to the no-slip condition to higher velocities in the free-stream (*57*). The boundary layer also allows small fishes or benthic species to use their fins to generate downwardly directed force and increase the frictional contact force with the substrate. This enables fish to hold position with minimal body movement even in high-velocity flows (*57*) (*58*) (*59*, *60*) (*61*), largely negating the hydrodynamic challenge from increased drag associated with living in rapid flows (*54*).

Although entrainment near obstacles, Kármán vortex locomotion, and boundary-layer station holding discussed here all exploit spatiotemporal variation in flow, the shear-layer gait represents an additional mechanism for harnessing energy from flows. Unlike fish exhibiting the Kármán gait in vortex flows (Fig. 9D) (*33*), which involves relatively large and mostly passive body deformation within the vortex street, the shear-layer gait is characterized by minimal undulation and near-static posture at an angle of attack within the shear layer (suppl. movies 1– 3). Unlike boundary-layer station holding, fish using the shear-layer gait are exposed to high free-stream velocities and do not maintain position by frictional contact, but by relying on a force balance within the velocity gradient (Fig. 2 & 6).

### Fish body posture, kinematics and energetics

Locomotion in a fluid environment is energetically costly, even when fish hover in near-zero velocity (*62*) (*63*) (*64*). Yet, the shear-layer gait displayed a total energy expenditure (aerobic and non-aerobic) that was indistinguishable from the resting level, despite exposing the entire lateral side and the head to freestream velocity equal to 50% of their maximum swimming speed, where their locomotor cost was 74% lower than free-stream swimming at the same velocity (Fig. 5). In comparison, entrainment behaviours have an approximately 36% reduction in aerobic energetic compared to free-stream swimming in trout at a similar speed (1.8 BLs^-1^) to that studied here (1.5 BLs^-1^), and the energy saving can reach up to 62% at 3.5 BLs^-1^ when fish entrain by a D-cylinder (*51*). Trout swimming laterally or behind a cylinder reduced their energetic expenditure by approximately 12–33% over 1–3 BLs^-1^ (*46*). Trout utilizing a Kármán gait at 3.5 BLs^-1^ downstream of a D-cylinder exhibit a 55% energy saving compared to the same freestream velocity (*51*). Hence, the shear-layer gait yields larger energetic savings than the Kármán gait and is nominally larger than the maximal energy savings reported for lateral entrainment, providing a particularly powerful energy-saving strategy.

The extent of energy saving within a shear layer reflects both reduced locomotor mechanical work and ventilatory costs. Trout within the shear layer, at certain times, nearly abolished undulation, and Amp_tail_ and *f*_tail_ approach zero (Fig. 2; suppl. movies 1-3). Fish in the shear layer can also passively ram ventilate, where water passes through the oral cavity and over the gills. This avoids active buccal pumping that can otherwise consume up to ∼15% of resting metabolic rate (*65*) (*66*) (*67*) (*68*), and explains how fish can use passive fluid mechanisms to assist in oxygen extraction when in a velocity gradient to reduce their energy cost so substantially.

Synchronized measurements of dynamic aerobic metabolic rate and kinematic modulation (*69*) (*70*) revealed a temporal coupling between kinematics and metabolic rate as individuals encountered the alternating conditions between the free stream and the shear layer. We showed that the transitions between a shear-layer gait and free-stream undulation are accompanied by rapid, tightly coupled changes in kinematics and metabolic rate. When the shear layer is removed, metabolic rate and tail undulatory movements increase rapidly; when the shear layer is restored, fish re-enter the velocity gradient, re-establish the shear-layer gait, and return to near resting metabolic rates (Fig. 5E & 5F). Synchronized measurements of dynamic metabolic rate and locomotor kinematics modulations are key to understanding the cost of moving under different hydrodynamic conditions (*70*).

To our knowledge, no previous study has explored energy-saving mechanisms where kinematic modulation is temporally coupled with dynamic changes in metabolic rate, and the conventional sequential measurements can underestimate the actual costs (Fig. 5E & 5F). This provides a novel and potentially powerful approach for future studies of metabolic energetic costs of various locomotion gaits in aerial, terrestrial and aquatic organisms (*70*) (*71*).

### The emergence of net-force equilibrium through fluid mechanics

Experimental measurements of flow fields and forces, combined with computational fluid dynamics (CFD), show that the energy-saving mechanisms of the shear-layer gait arise from an overall net-zero force balance achieved within the velocity gradient. These regions of low net force overlap with the locations selected by trout, indicating that fish position themselves within a zero-net-force corridor sculpted by the gradients of fluid velocity and pressure (Fig. 9A).

Three-dimensional CFD simulations based on the body kinematics reconstructed from high-speed videography illustrate the mechanisms of force balance (Fig. 7). As water flows around the foil, a broad region of reduced pressure extends laterally from the leading edge and the wake behind (Fig. 7B, C, D; Fig. 9). When a fish station holds in the shear layer, ipsilateral side of the body encounters near free-stream velocity (Fig. 3,7; suppl. movies 3a, b, c) and experiences near constant viscous drag (Fig. 7C), while the opposite side encounters reduced velocity and pressure, generating anteriorly directed suction forces. Pressure reduction behind the foil generates forward pressure on the fish body surface, while modest oscillations are present (Fig. 7E, F, D). On average, the negative (forward) pressure forces balance the positive (backward) viscous drag on the body, while the low-amplitude undulation supplies only momentary thrust to maintain position (Fig. 7B). The total drag coefficient (viscous and pressure) and tail thrust both oscillate around zero. This contrasts with fish swimming in uniform flow, where the area immediately in front of the head of swimming fish can experience a positive stagnation pressure (Fig. 7D) (*72*), resulting in increased drag.

Crucially, trout frequently choose to occupy the steep velocity gradient, rather than the low-velocity core directly downstream of the foil (Fig. 9A). The wake zone is characterized by large, unstable lateral velocity fluctuations that induce greater postural instability (Fig. 8B,C). In contrast, shear-layer regions provide a broader and more stable pressure field and a restoring force condition. Small medial and lateral deviations in location could tend to return the fish toward the equilibrium corridor, rather than displacing it further into the free stream or deep into the wake (suppl. movie 6). These stabilizing fluid conditions enable fish to sustain the shear-layer gait with minimal corrective motion.

Two-dimensional CFD analyses across a wide range of foil angles (0–90°) showed that time-averaged drag on a fish-like body transitions from positive (drag) to negative (thrust) as foil angle increases (Fig. 8, suppl. movie 5), where fluctuations of drag and lift grow with increasing angle and shear-layer strength. These simulations, together with experiments, illustrated that maximum energetic benefits occur at intermediate shear-layer strengths (30°**–**45° foil angles; average shear layer strength: 0.044, 0.037 m·s^-1^·mm^-1^; coefficient of variation: 10.7, 26.5%; Figs. 3, 4, 8). Where the time-averaged drag coefficient is centred around zero, force fluctuations are reduced, and kinematic effort is the lowest. At higher foil angles (90°), the stronger and more variable shear layers lead to increased kinematic response in live fish and greater spontaneous force fluctuations in model fish.

### Fluid-mediated locomotor energy saving

Altogether, posture and orientation of the fish’s foil-like body act as a passive fluid stabilizer that interacts with the velocity gradient to achieve net force equilibrium without continuous and active propulsion. This extends classical views about propulsion enhancement and drag reduction (*73*) (*73*) by revealing how a flexible body can exploit the spatial structure of shear flows to generate passive thrust. The shear-layer gait and its underlying force balance persist across a range of velocity gradients, although kinematic and energetic dampening is most effective at intermediate shear-layer strength. Thus, using a shear-layer locomotor strategy may be available to fishes in diverse natural flow geometries, including wakes of rocks, logs, vegetation, conspecifics, and around artificial structures (*see* suppl. movie 7 for a coho salmon using a rock with an angled surface to locate within a shear layer in a stream).

Nevertheless, several open questions remain. First, how do different species, body shapes, and size classes employ shear-layer strategies in rivers, coastal zones, and possibly pelagic environments? Second, how do fine-scale fin movements interact with oscillatory fluid forces to assist in maintaining position within the shear layer? Fish may exhibit compensatory fin movements to station in shear layers and fine-tune body position (suppl. movie 1; Fig. 3,7).

Third, how do sensory systems, particularly flow sensors and visual inputs, guide precise body positioning and the postural modulations needed to utilize the zero-net-force corridor? Fourth, how frequently do other animal taxa exploit such shear-layer energy harvesting during migration, foraging, or predator-prey interactions, and does this shape life-history and ecological niches?

For example, the wing morphology of birds is linked with the kinematic modulation of stability control (*74*) (*75*), and the covert feathers on the lower surface of bird wings utilize shear layers to enhance lift and reduce drag for flight (*76*).

The understanding of a causal connection from fluid dynamics to kinematics and energetics provides a framework that may be extended to aerial and terrestrial systems where shear layers and sharp velocity gradients are also pervasive. Advancing our understanding of animal use of fluid velocity gradients will require an integration of field studies, tagging and tracking animal movement patterns, and controlled laboratory experiments coupled with further advances in experimental and computational fluid dynamics.

## CONCLUSION

Animals and vehicles moving through fluids routinely encounter a variety of fluid dynamic challenges that include changes in pressure, velocity, temperature and viscosity. For example, fish migrating upstream (*77*) (*78*) (*79*) (*80*), insects and birds navigating around obstacles (*81*) (*82*) (*83*), deep-diving marine mammals (*84*) (*85*) (*86*), and birds and insects (*87*) flying at altitude and in cluttered environments encounter changes in fluid velocity (*88*) (*83*). These are all conditions that challenge the locomotor capabilities of animals. More broadly, movement is energetically expensive, especially in the dense and viscous aquatic medium where drag scales exponentially with speed. Active propulsion through challenging hydrodynamic conditions, where strong velocity gradients are abundant, can be taxing for the energetic capacity of animals (*89*) (*90*) and for the energy storage capacity of vehicles.

By integrating behavioural quantifications with analyses of kinematics, energetics, and experimental and computational fluid mechanics, our results demonstrate that fish can exploit fluid velocity gradients for efficient movement by passively generating thrust using the external fluid kinetic energy. This establishes a mechanistic framework for understanding passive thrust generation and force equilibrium in fluid velocity gradients, which connects the physics of shear layers to energetic strategies of animals, and contributes to understanding how diverse environmental flow conditions can influence organismal ecology and evolutionary fitness.

Velocity gradients are not merely a locomotor challenge. They also represent opportunities for energy harvesting by animals. The principles uncovered in fish interacting with shear layers have direct relevance to aerial locomotion in birds and insects, and offer a design blueprint for autonomous underwater (and aerial) vehicles and flow-sensing robots. All these systems need to navigate through complex, unsteady and heterogeneous environments, and can benefit from not resisting ambient flow gradients, but instead by becoming a part of the flow via strategic positioning within fluid velocity gradients to harness kinetic energy and minimize power consumption.

## MATERIALS AND METHODS

### Experimental animals

We used brook trout (*Salvelinus fontinalis*) that were acquired from a local hatchery near Boston, Massachusetts USA, as our study organism. Fish were housed in a tank (1000 L) with self-contained thermal control (16 °C), an aeration system (>95 % air saturation) and a filtration system (Fluval FX6, Rolf C. Hagen Corp. Italy). The filtration system was routinely cleaned.

Water changes and water quality monitoring were routinely carried out by a designated technician. Fish were fed *ad libitum* three times weekly (Keystone Pellets, Skretting, USA). Animal holding and experimental procedures were approved by the Harvard Animal Care IACUC Committee (protocol number 20-03-3). Three brook trout (Fork length: 22.5 – 24 cm mass: 207-272g) were tested for integrated bioenergetics and biomechanics tests, and five trout similarly sized trout (21– 24cm) were used for behaviour, kinematics and hydrodynamics experiments.

### Experimental arenas

A series of shear layer flow fields was generated using a NACA 0012 foil (span: 105 mm; cord: 67 mm; thickness: 8.1 mm; center of rotation: 48 mm from trailing edge; material: transparent photopolymer [RGD810] from a Connex 500 3D printer). Fish swimming in freestream flow occurred with the foil rotated to a 0° angle to oncoming flow, and shear layers lateral to each foil edge were created by orienting the foil at 30°, 45° or 90° to streamwise flow.

We used two recirculating flow tanks for the experiments reported here. First, analyses of metabolic energy expenditure (*see* below) were obtained simultaneously with trout body kinematics. We used two synchronized lateral and ventral high-speed Promon U1000 high-speed cameras (AOS Technologies Switzerland) to image the central region of the respirometer as in previous research (*63*). The details of this system are given in the supplementary materials. The core of the system was a customized swimming tunnel respirometer (Loligo Systems, Denmark; 46 L total volume; working section 142 × 142 × 500 mm) that allowed measurement of oxygen consumption (*see below*) and clear sections that enabled video recording from the lateral and ventral views (*63*) (*64*) (*91*) (*92*). We modified the respirometer to include a rotating foil with a sealed shaft that allows dynamic changes in the flow regime to be experienced by trout within the respirometer. Flow velocity patterns within the central respirometer chamber were calibrated using particle image velocimetry.

Second, additional trout behaviour, body kinematics, and hydrodynamics experiments were conducted in a second flow tank. This tank had a working section 28 cm × 28 cm × 64 cm (total volume = 570L). The front and back portions of the swimming section were separated by honeycomb flow regulators, and videos were recorded from the ventral and lateral perspectives. Particle image velocimetry was used to measure flow patterns in the working section (*see* below) and to examine flow around the foil both with and without a trout present. This flow tank also contained a robotic mechanism (*93*) to control the angle and position of a hydrofoil in the flow and enabled an integration of a force sensor (*see* below) to measure forces in relation to a foil oriented at angles of 0°, 30°, 45°, and 90° at freestream flow velocity of 0.45 m·s^-1^, 50% of the maximum sustained swimming speed of the trout tested (Fig. 6 A,B).

Water velocity in the free stream for all tests varied from 1.5 – 2.0 body lengths s^-1^ (BL s^-1^) during swimming (giving Reynolds numbers on the order of 7•10^4^ – 9 •10^4^, using trout body length as the characteristic length) for trout located in the free stream.

### Kinematic analyses

Midline kinematic extraction involved measuring body kinematics sampled evenly across 11 time points within each tail beat cycle and included the extreme amplitudes of each tail beat cycle. The midline started at the tip of the nose and ended at the tip of the caudal fin and was quantified for trout swimming both in the free stream with the foil oriented at °0, and also during shear layer locomotion occurring at foil angles of 30°, 45°, and 90°. Mid-line characterization used previously published custom code (CurveMapper, (*94*)) in MATLAB (v.2024) (MathWorks Inc., Natick, MA, USA). CurveMapper fitted a spline curve equation using 200 equally spaced coordinates on the selected points along the midline of the fish (*e.g.*, tip of snout = point 0; tip of caudal fin = point 200). The absolute displacement based on the midline of the fish was then compared across different foil angles.

Tail beat frequency (*f*_tail_) and tail beat amplitude (<u>Amp_tail_</u>) were tracked by artificial-intelligence-assisted computer vision functions in DLTdv8 (*95*) (n=7-10). Analysis of the values for *f*_tail_ was conducted in Labchart (v8.1.3, ADInstruments, Dunedin, New Zealand), using the sine wave algorithm in the frequency measurement function. <u>Amp_tail_</u> was extracted by calculating the absolute differences between peaks and valleys of the sine wave within a tail beat cycle.

Characterization of shear layer locomotion, when fish moved between freestream flow and returned to the shear layer, was also analyzed by artificial-intelligence-assisted computer vision functions in DLTdv8 (*95*). Tracking of tail movement was done continuously for ∼40 mins, resulting in ∼144,000 total frames from which frequency and amplitude were extracted. Tail movement data were analyzed by Labchart (v8.1.3, ADInstruments, Dunedin, New Zealand) to calculate a continuous measurement of *f*_tail_. Video recordings were synchronized to the measurements of dynamic metabolic rate.

### Behavioural heat map analyses

The effects of the shear layer flow field on the two-dimensional spatial location of the fish were studied by a series of behavioural trials. A fish was acclimated to the flow tank environment for at least 1 hour before each trial. Flow velocity was then set to 0.45 m·s^-1,^ and trout were subsequently exposed to the foil at 0°, 45°, and 90°, for 10 minutes each. The order of fish exposure to each foil angle was randomized. Throughout this 30-minute period, video from the ventral view was recorded (n=5 fishes) using a Promon Y1000 high-speed camera (AOS Technologies Switzerland) at 125 FPS.

To quantify how fish use different areas of the flow field, we tracked the fish’s position using DeepLabCut (*96*) at 1Hz. The neural network was trained on an initial 20 frames, and retraining and manual labeling was iteratively conducted until tracking was visually accurate. The distribution in location for each fish and each foil condition was estimated using a 2D kernel density estimator, with a 5 cm bandwidth. The ‘core area use’ was calculated as the minimum area that contained 50% of each fishes density (i.e. ∼ smallest area where fish spent 50% of their time). The orientation of the fish towards each side of the foil was calculated as the angle between the head and anal fin of the fish. The statistical and tracking analyses of the behavioural heat map were conducted in R version 4.4.2 (R Core Team).

### Experimental hydrodynamics visualization and analysis

To quantify the fluid dynamics underpinning the shear-layer gait, we used particle image velocimetry as established previously (*93*) (*72*). Particle image velocimetry was conducted to quantify flow patterns when the foil was oriented at angles of 0^◦^, 30^◦^, 45^◦^, and 90^◦^ to measure shear layer strength and the flow regime without fish. A horizontal plane of laser sheet was generated using a solid-state 532nm green laser (LD solid-state green laser, 5W, MGL-N-532A, Opto Engine LLC) that allowed visualization of flow around the foil and fish body (Fig. 4). Flow velocity was calculated from sequential high-speed video frames video frames (Photron AX50 high-speed camera, 1024×1024 pixels, 1.5 long second video, framerate = 1000 FPS, resolution = 1280 x1024) using DaVis v8.3.1 (LaVision Inc., Göttingen, Germany). A vector field (covering a horizontal plane of 34.3 cm^2^ with 2193 vectors) was characterized by a sequential cross-correlation algorithm applied with an initial interrogation window size of 64 × 64 pixels that ended at 12 × 12 pixels (3 passes, overlap 50%). OpenPIV-Matlab was also used to prepare flow visualizations, as in supplemental movie 3c.

We also quantified the hydrodynamic conditions when fish were swimming in both laminar and shear layer conditions under the same freestream velocity (1.5 BL s^-1^, ∼36.7 cm s^-1^) with the foil oriented at angles of 0^◦^, 30^◦^, 45^◦^, and 90^◦^. The fluid velocity field was calculated from consecutive video frames (1984 × 1032 pixels) using DaVis v8.3.1 (LaVision Inc., Göttingen, Germany). A vector field (covering a horizontal plane of 765.3 cm^2^ with 7998 vectors for calculations) was characterized by a sequential cross-correlation algorithm applied with an initial interrogation window size of 32 × 32 pixels that ended at 16 × 16 pixels with 3 passes at each window size that overlap 50%. From the fluid field, we calculated the streamwise velocity across cross-sectional profiles of frontal (F), head (H), pectoral (P), trunk (TK), tail (T) and post caudal (PC) locations as shown in Fig. 3.

We conducted two series of fluid dynamics analyses on the shear layer: (1) characterizing the velocity gradients across the swimming section, and (2) quantifying the strength of the shear layer. We quantified flow velocities in the experimental test arenas both when trout were present and holding position in the shear layer and in the vortex wake of the foil alone. We characterized the velocity gradient on the selected regions (see Fig. 3B) of frontal, head, pectoral, trunk, tail and post-caudal when fish were holding stationary in the shear layer. To demonstrate the effects of fish presence in the shear layer, we characterized the velocity gradients at the frontal and post-caudal regions in the flow field without the fish. To standardize the comparisons of the shear layer strength across different laboratory and field-testing conditions, we calculated the rate of change in velocity across the shear layer region (a proxy for shear layer strength). We standardized the sampling region as a 55-mm vertical line from the centre of the foil and ∼100 mm downstream from the centre of the foil (see Fig. 4A).

### Dynamic and steady-state locomotor energetics

We quantified locomotor energetics when trout swim in the flow field created by the 30° foil angle and compared energy use to free-stream locomotion. We used an established Integrated Biomechanics & Bioenergetic Assessment System (IBAS) (suppl. materials; (*64*)(*63*)) to quantify the energetics and time-matched kinematics of shear layer locomotion using the 30° foil to improve the fish biomass to flow tank volume ratio and ensure a strong signal-to-noise ratio for the measurements of metabolic rate. When testing the 45° foil, the shear layer extended too close to the wall of the swim-tunnel respirometer, which introduced a wall effect as a confounding factor. Thus, we used the 30° foil to create shear layer located at the center of the flow tank to avoid wall effects.

Swimming performance test trials were conducted with *Salvelinus fontinalis* fasted for at least 48 hours to reach a post-absorptive state (*97*) (*98*). Before the swimming performance test, the testing fish were weighed and placed in the swim-tunnel respirometer. The fish swam at ∼30% *U*_crit_ for ∼30 mins to help oxidize the inevitable but minor lactate accumulation during the prior handling and help fish become accustomed to the flow conditions in the swim-tunnel respirometer (*99*). After initial habituation, the fish was rested for more than 20 hours in the respirometer under quiescent and undisturbed conditions. During this time, we used an automatic system to measure the resting *Ṁ*O_2_ for at least 20 hours. Software (AquaResp v.3, Denmark) and relays (Cleware GmbH, Schleswig, Germany) were used to control the intermittent flushing of the respirometer with fresh water throughout the trial to ensure a high O_2_ saturation of the respirometer water. *Ṁ*O_2_ was calculated from the continuously recorded dissolved O_2_ level (at 1 Hz) through a homogenous water sampling loop to measure the decline of water dissolved O_2_ due to animal respiration. The intermittent flow of aerated water into the respirometer occurred over a cycle of 960 s, with 120 s of the aerated water being introduced into the respirometer and 840 s where the pump was off, and the respirometer became a closed system. The first 180 s after each time the flushing pump was turned off were not used to measure *Ṁ*O_2_ to allow O_2_ levels inside the respirometer to stabilize. The remaining 660 s when the pumps were off during the cycle were used to measure *Ṁ*O_2_. The in-line pump for water homogenous loop for O_2_ measurement was constantly on over the entire experimental trial.

We studied three individual trout, and each individual was measured with a repeated design that generated six instances of metabolic rate measurement (Eqn. 1 in suppl. materials) to quantify the energetic costs of each locomotion gait and during the transitions between swimming in the foil shear layer and sustained undulatory swimming gait in the free stream.

In addition, we measured steady-state locomotion when fish were swimming for a prolonged time within the shear layer and separately when fish swam for extended periods in freestream laminar flow. At each fluid condition, fish swam for 12 mins (*100*) (*101*) to reach a stable aerobic metabolic rate (analyzed using Eqn 2 in suppl. materials). These results were included in the steady-state metabolic rate analyses as shown in Fig. 5 A&B.

The goal of the steady-state locomotion protocol was to measure the contribution of non-aerobic O_2_ cost, where most of the non-aerobic cost for trout swimming at 1.5 BL s^-1^ was related to replenishing high-energy phosphate and the O_2_ stores in haemoglobin and myoglobin (*102*). Measuring the non-aerobic cost together with the aerobic cost in the same tests for the same individuals enables calculation of the total energy expenditure (*63*) (*64*). We measured excess post-exercise oxygen consumption (EPOC) after the *U*_crit_ test for the ensuing ∼1 hour, recorded by the respirometer system described above. From our experimental traces, nearly all fish returned to the pre-testing resting metabolic rate within ∼15 mins. To best capture the rapid decline of metabolic rate at the start of EPOC, the respirometer remained closed after sustained locomotion for ∼25 mins (*see* Eqn 2). Upon the completion of the two-day protocol, the fish were returned to their home aquarium for recovery. Fish condition was closely monitored during the first ∼36 hours after the experiment, during which no mortality was observed.

Experimental shear layers for fish locomotion were created using a NACA 0012 foil (span: 105 mm; cord: 67 mm; thickness: 8.1 mm; center of rotation: 48 mm from trailing edge; material: transparent photopolymer [RGD810] from an Connex 500 3D printer) oriented at a 30-degree angle to free stream flow and positioned within the swimming section. The transparent photopolymer reduced the potential visual effects of a solid object in front of the fish, while allowing the fish to locate the shear layer region generated by the 30° foil through flow sensing. The foil was connected to a shaft sealed through a hole drilled through the surface of the respirometer. The drilled hole was tapped, fitted with a watertight cable gland, and sealed with epoxy. A watertight cable gland enabled the foil to be rotated between 30° and 0° to streamwise flow while the fish was swimming. This arrangement allowed for the foil angle to be altered by rotating the shaft without opening the respirometer and for fish to transition between shear layer locomotion and free stream undulatory locomotion, and back again, while metabolic rates were recorded.

### Force measurement

We measured the forces acting on a rigid model fish to understand the fluid forces acting on the fish body and how these forces were spatially variable (Fig. 6). We used a 3D model of a trout that was previously used in computational fluid dynamics studies (suppl. materials, Trout.model.stl). This model was 3D printed in Nylon PA12, a rigid black material. The model was mounted to a rod vertically from its center of mass. We used a trout model 25 cm long, which corresponded to the size of the fish used in our study.

This model fish was suspended from the robotic mechanism in the kinematic swimming tunnel. The model fish’s lateral position (stream width, Y axis) and yaw angle to oncoming flow were digitally controlled by a computer. Its streamwise position (X axis) was controlled by the experimenter. A six-axis force and torque sensor (ATI Nano-17, Apex North Carolina) was mounted to the rod that connects the model fish and the robotic mechanism, which estimated the forces and torques acting on the model fish.

We fixed a NACA 0012 foil at either 45° or 90° to the oncoming flow and set the water velocity in the flow tank to 0.45 m·s^-1^. The fish model was then sequentially placed in ∼100 different locations downstream and lateral to the foil. These locations formed a grid (112 measurement points), with the streamwise resolution of 3cm, and a stream width resolution of 1cm. At each location, the model was oriented at two yaw angles to the free stream flow (, 0°, and 10°). For each combination of location and model angle, the 3D forces and torques were recorded (at 1000 Hz) for 10 seconds using a custom LabView program.

For each sampling condition (model location and angle), the forces and torques were averaged over the entire 10-second period, and forces for the X and Y axes were rotated from the local fish frame to the global frame (X is streamwise, and Y is stream width), to account for the varying yaw angle of the fish model. The X, Y forces and the Z torque point estimates were spatially interpolated to create a continuous estimate of the horizontal plane map for the variations in force and torques (Fig. 6). Areas considered to be “low force” zones had to experience less that 30 millinewtons of force.

### Computational fluid dynamics

We conducted both two (2D) and three-dimensional (3D) computational fluid dynamics (CFD) simulations to understand forces on the body of the fish and in the vortex of the foil. A direct numerical simulation (DNS) algorithm was used to reach 2^nd^-order accuracy. In both 2D and 3D, the computational domain was discretized with Cartesian grids, on which the fish and foil geometries were resolved using the immersed-boundary method (IBM). The details of the numerical solver were described by Zhang et al. (*103*). The numerical solver has also been validated with both hydrofoil and fish experiments (*104*, *105*).

In the 3D simulation, the fish kinematics and movements were reconstructed from video recordings of a period when a fish held steady station with minor adjustments near the foil’s leading-edge side. The angle of attack for time-series analyses (snout-tail-base line relative to the incoming flow) was 4.2 degrees (range from −1.5 to 7.5 degrees). The average heading angle was slightly higher at 5.9 degrees (Range: 2.1 to 8.9 degrees). The Reynolds number in the simulation was set to 60000, with sub-grid structures resolved using dynamic Lagrangian turbulent modelling. The effect of the foil starting vortices on the forces acting on the body was mitigated by holding the fish static for a period and implementing reconstructed motion after the forces acting on the static fish model reached a steady state.

To understand the effect of foil angle on the forces acting on the fish, we conducted a series of 2D CFD simulations. An inclined, static NACA foil was employed to model a fish swimming downstream of a foil, as illustrated in the schematic in Fig. 8A. The foil leading edge to fish model leading edge distance is 0.9 foil chord length, and the lateral distance is 0.4 chord length. The fish model was oriented at a 10° angle of attack, a position and posture consistent with the shear-layer gait of biological fish. The Reynolds number was set to 5,000, based on the incoming flow velocity *U*_∞_ and the foil chord length *c*. A two-layer mesh refinement was used, with the finest grid size of Δmin = 3.25 × 10^-3^*c*. This resolution has been validated in previous studies (*106*). The simulations were performed over a duration of 60*T_c_*, where *T_c_* = *c* / *U_∞_* was the convective time scale based on the foil chord length, to ensure that the strongly nonlinear and highly unsteady flow at high angles of attack was fully developed and that statistically steady results were obtained. The time-averaged velocity is normalized as *U*∗ = (*U* − *U*)⁄*U*, where *U* is the time-averaged velocity and *U* is the incoming velocity. The variations in force amplitude were characterized by the root-mean-square (RMS) values of drag (C_D_) and lift (C_L_), and their time derivatives. Ċ_D_ and Ċ_L_, were also calculated to quantify how rapidly the force varies in time.

### Statistical analyses

Measurement points are presented as violin plots, where dashed lines are medians and dotted lines were quantiles, and the shape of the plot represents the frequency distribution. The regression models are presented as the mean ± 95% confidence interval. For the metrics that failed normality tests, square or logarithm transformations were applied to meet the assumptions of normality of residuals, homoscedasticity of the residuals, and no trend in the explanatory variables.

The statistical comparisons of kinematics and energetics used the Generalized Linear Mixed Effects Model with Holm–Šídák *post-hoc* tests. Fixed factors were 90°, 45° & 30° foil angles (three shear layer conditions) and 0° foil angles (freestream). The random effects were the assignment of fish to the flow field treatment. Statistical comparisons of two-dimensional CFD metrics used the General Linear Model with Holm–Šídák *post hoc* tests. Statistical significance was denoted by *, **, ***, **** for *p*-values of ≤ 0.05, ≤ 0.01, ≤ 0.001, ≤ 0.0001 respectively. These statistical analyses were conducted in SPSS v.28 (SPSS Inc. Chicago, IL, USA) or GraphPad, Prism v.10 (GraphPad Software, LLC, Boston, MA, USA).

Analyses of fish movement area size and angle (Fig. 1) were conducted in base R (*107*). Statistical graphics were generated in GraphPad, Prism v.10 (GraphPad Software, LLC, Boston, MA, USA).

## Acknowledgments

Many thanks to members of Lauder Laboratory for numerous discussions about fish locomotor behaviour, for comments on the manuscript, to Cory Hahn for fish care, and to K. Liebich from U.S. Fish & Wildlife Service for providing the wild fish video

## Author contributions

Conceptualization: YZ, CW, RT, GL

Methodology: YZ, GL, CW, RT, JG, YP, HD

Investigation: YZ, CW, RT, JG, YP, HD

Visualization: YZ, CW, RT, JG, YP, HD

Funding acquisition: GL, YZ

Project administration: GL, YZ, CW, RT

Supervision: GL

Writing – original draft: YZ

Writing – review & editing: YZ, GL, CW, RT, JG, YP, HD

## Data and materials availability

All data are available in the main text or the supplementary materials.

## Supplementary Materials

### Supplementary Movies, S1 to S7

**Movie S1. Fish transition from shear layer flow to freestream flow conditions.** A light movie that demonstrates brook trout (*Salvelinus fontinalis*) exhibiting dampened undulatory motion in the shear layer created by a NACA 0012 foil angled at 45°. Once the foil is turned to 0°, the trout starts to exhibit typical undulatory locomotor movements in freestream flow conditions.

**Movie S2. Fish exhibiting a steady shear-layer gait.** Movie that demonstrates a brook trout (*Salvelinus fontinalis*) holding station in the shear layer created by the NACA 0012 foil angled at 30°. The trout shows almost no undulatory motion.

**Movie S3. Particle image velocimetry (PIV) of fish exhibiting the shear-layer gait.** (a) High-speed PIV video showing a brook trout (*Salvelinus fontinalis*) exhibiting the shear-layer gait. Fluid flow is visualized by the particles illuminated by two laser sheets (both sides of the lateral view) to remove shadows. (b) PIV video showing the trout and the shed vortices (visualized by vorticity shown in red and blue colours) to illustrate the fluid dynamic mechanisms underlying the shear-layer gait. (c) PIV footage of velocity and vorticity during fish locomotion in the shear layer. The high and low flow regions on both sides of the fish create a fluid pressure gradient, which acts on the solid surface of the fish body and results in a force equilibrium, allowing for station holding behind the foil.

**Movie S4. Three-dimensional computational fluid dynamics (CFD) analysis of the shear-layer gait.** (a) Frontal and (b) perspective views of the CFD visualization of trout exhibiting a shear-layer gait in the fluid velocity gradient. Fish locomotion is based on videos of trout swimming in the fluid velocity gradient. The NACA 0012 foil was angled at 45° in the CFD analyses.

**Movie S5. Two-dimensional computational fluid dynamics (CFD) analysis of flow shed by a foil relative to a fish body model in the position of the shear-layer gait.** Video shows vortices shed by the foil through time and how they impact the fish body model. Figure 8 in the manuscript shows the data from this simulation when the foil is oriented at a range of angles.

**Movie S6. Animation showing the force balance on a trout body model.** Black dots indicate the location of each force measurement on a trout body model (not shown) with respect to a foil oriented at a 45 degree angle. Yellow arrows show the direction of net force in the streamwise and lateral directions. Animation shows the convergence of upstream force into the low velocity corridor downstream of the foil.

**Movie S7. Wild coho salmon (*Oncorhynchus kisutch*) exhibited a shear-layer gait lateral to a sloped rock and in fast-moving flow.** The motions of aquatic vegetation and bubbles show that the fish is oriented against the fast-moving flow. The red coho salmon positioned behind a sloped rock had a low and nearly absent undulatory motion in comparison to other fish in the background. The video was taken in Campbell Creek, Anchorage, Alaska. (Credit: K. Liebich, U.S. Fish & Wildlife Service).

**Data S1. Three-dimensional body model of a trout (.stl).** This trout model was 3D printed and attached to a six-axis force/torque sensor to map the forces generated in the wake of a foil. Data from these experiments are shown in Figure 6 of the manuscript.

## SUPPLEMENTARY TEXT

### Experimental system – Integrated Biomechanics & Bioenergetic Assessment System (IBAS)

The core of our experimental system for quantifying fish energy use is a 46-l (respirometry volume plus tubing) customized Loligo® swim-tunnel respirometer (Tjele, Denmark). The respirometer was equipped with an electric motor that drove a shaft attached to a propeller sealed inside the respirometer. The water velocity of the respirometer was controlled by regulating the revolutions per minute (RPM) of the motor. The detailed descriptions of the experimental systems are in the previous publications, and here we documented the systematic parameters specific to this study (*1*)(*2*).

We specifically designed the swim-tunnel respirometer as oval-shaped. Incorporating A hollow space in the centre of the respirometry increases the turning radius of the water current. As a result, the water velocity passing the swimming section (cross-section: 142 × 142 × 500 mm) is more homogeneous (validated by PIV). To create the controlled flow condition, a honeycomb flow straightener (142 × 142 × 155 mm) is installed upstream of the swimming section to create laminar flow (validated by PIV). The linear regression equation between RPM and water velocity (V, cm s^-1^) of laminar flow is established (V = 0.1317 • RPM – 6.259, R^2^ = 0.9995, *p <* 0.0001, Fig. S1) by the velocity field measured by particle image velocimetry (PIV). The free-stream laminar flow is set as 1.5 BL s^-1^, which is in the RPM (revolutions per minute) range of 309 – 350.

The key to measuring the dynamic aerobic rate is to have the instrument achieve a higher-than-standard signal-to-noise ratio for the measurement of dissolved O_2_ in water. The rate of dissolved O_2_ being consumed by the animals inside the respirometer measures the whole-organism metabolic rate. To facilitate this goal, we specially designed a water homogenous loop is installed 125 mm downstream of the propeller and the water is returned to the respirometer 420 mm before the swimming section. To avoid mixing the counter currents, the water in the homogenous loop flowed (designated in-line circulation pump, Universal 600, EHEIM GmbH & Co KG, Deizisau, Germany) in the same direction as the water flow in the swimming tunnel. The dissolved O_2_ level in the water was continuously measured by high-resolution fibre optic O_2_ probe (recorded at a frequency of ∼1 Hz & response time < 15s; Robust oxygen probe OXROB2, PyroScience GmbH, Aachen, Germany) that was sealed in the homogenous loop at downstream of the circulation pump to better homogenize the dissolved O_2_ gradient of the sampled water.

The oxygen probe was calibrated to 0 % sat. (a solution created by super-saturated sodium sulphite and bubbling nitrogen gas) and 100 % sat. (fully aerated water). The pre-filtered water (laboratory grade filtration system) is constantly disinfected by six units of UV light (6 units of JUP-01, SunSun, China) located in an external water reservoir to suppress bacterial growth. The background *Ṁ*O_2_ in the swim-tunnel respirometer was measured for ∼30 min before and after each trial. The average background *Ṁ*O_2_ (< 0.5% of fish *Ṁ*O_2_) was negligible for *Ṁ*O_2_ correction. Water changes of ∼70% total volume occurred every other day and a weekly disinfection is conducted using sodium hypochlorite (Performance bleach, Clorox & 1000 ppm).

The integrated study of bioenergetic and biomechanics was achieved by simultaneously measuring three-dimensional swimming kinematics. In this regard, the customized oval-shaped swim-tunnel respirometer has a clear side panel in the swimming section that provides a lateral view. To obtain a ventral view, the swimming section also has a clear bottom panel and the entire swim-tunnel respirometer locates on a platform (elevated 295 mm above the base) with an open window beneath the swimming section and a front surface mirror to be installed at a 45° angle underneath the swimming section. The three-dimensional reconstruction of the fish kinematics using the time-matched high-speed videography recorded in two streaming cameras (PROMON U1000, AOS Technologies AG, Switzerland, lens: Nikon AF-S NIKKOR 17-35 1:2.8D, Japan). The lateral-view camera is positioned 920 mm to the side of the swimming section, and the ventral-view camera is positioned 670 mm to the side of the swimming section. Synchronized lateral and ventral video recordings were recorded at 60 fps (exposure, lateral view: 10648 us, ventral view: 12099), and each frame was 1920 by 687 pixels (ventral view) and 1952 by 575 pixels (lateral view). To avoid light refraction passing through the water and distorting the video recordings, the swim-tunnel respirometry is not submerged in the water bath like the regular swim-tunnel respirometry. Temperature regulation of the respirometer is achieved by regulating room temperature with climate control units, installing thermal insulation layers on the respirometry and replenishing the water inside the respirometer from a thermally regulated (chiller: ActiveAQUA Water Cooling System, Hydrofarm, USA) water reservoir (thermal insulated 156-L aquarium) located externally.

The introduction of the aerated water (100% sat., air system built into the laboratory of the Museum of Comparative Zoology) to the respirometer uses an in-line pump (Universal 3400, EHEIM GmbH & Co KG, Deizisau, Germany) and an in-line computer-controlled motorized ball valve (U.S. Solid) installed at the in-flow tube. The outflow tube is also equipped with a one-way valve and a manual valve. As a precautionary practice to eliminate the exchange of water between the respirometer and the external reservoir when the water moves at a velocity inside the respirometer, the manual valve is shut during the measurement period. The flushing was manually controlled to maintain dissolved O_2_ above 80 % sat. When the respirometer was closed to measure *Ṁ*O_2_, the water temperature fluctuated no more than ∼0.2 °C (measured by a needle temperature probe, Shielded dipping probe, PyroScience GmbH, Aachen, Germany, sealed through a tight rubber port of the respirometer).

To allow fish to reach a quiescent state before the experimental trial, the foot traffic around the experimental platform is restrained to the absolute minimum. Fish were oriented by small, dual anterior spots of white light (lowest light intensity, Model 1177, Cambridge Instruments Inc, New York, United States) for orientation (one to the top and the other to the side) of the swimming section. The entire Integrated Biomechanics & Bioenergetic Assessment Platform (IBAP) is covered by a laser blackout sheet (Nylon Fabric with Polyurethane Coating; Thorlabs Inc, New Jersey, United States). The test section is illuminated by infrared light arrays that do not introduce visual distraction to the fish.

### Bioenergetic measurement and modelling

To estimate the steady-rate whole-animal aerobic metabolic rate, *Ṁ*O_2_ values were calculated from the sequential interval regression algorithm (Eqn. 1) using the dissolved O_2_ (DO) points continuously sampled (∼1 Hz) from the respirometer. The estimate of dynamic aerobic metabolic rate uses a rolling regression algorithm (Eqn. 2)

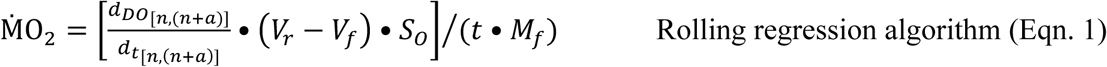

Rolling regression algorithm (Eqn. 1)

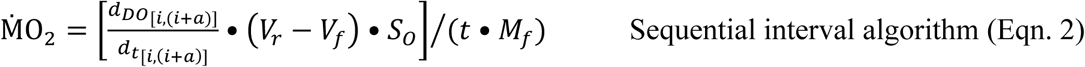

Sequential interval algorithm (Eqn. 2)

Where units for *Ṁ*O_2_ are mg O_2_ h^-1^kg^-1^, *d_Do_*/*d_t_* is the change in O_2_ saturation with time, *V_r_* is the respirometer volume, *V_f_* is the fish volume (1 g body mass = 1 ml water), *S_o_* is the water solubility of O_2_ (calculated by AquaResp v.3 software) at the experimental temperature, salinity and atmospheric pressure, *t* is a time constant of 3600 s h^-1^, *M_f_* is fish mass, and *a* is the sampling window duration, *n* is 1 DO increment from the first DO measurement at a set sampling frequency of 1 Hz, and *i* is the next DO sample after the preceding sampling window. *Ṁ*O_2_ of the low data quality (*e.g.* non-linear decline or R^2^ <0.5) were removed to control for the measurement errors (*3*) (*4*) (*5*).

The resting oxygen uptake (*Ṁ*O_2rest_), estimating the resting metabolic demands of an individual, is calculated from a quantile 20% algorithm (*4*) using the *Ṁ*O_2_ estimated between the 3^rd^ to the ∼24^th^ hours after the swimming test. The algorithm removed the metabolic costs of spontaneous activity to achieve a reliable estimate of the resting metabolic demands.

The excess post-exercise O_2_ consumption (EPOC) is an integral area of *Ṁ*O_2_ measured during post-exercise recovery, from the end of the locomotion activity until it reached *Ṁ*O_2rest_ plus 10% (*6*). This approach reduces the likelihood of overestimating EPOC due to spontaneous activities (*6*), while estimating the anaerobic costs for activities (*7*) (*8*) (*9*) (*10*).

We sum EPOC (*i.e.* non-aerobic O_2_ cost) with the aerobic costs over 12-min duration of locomotion to estimate a *total* O_2_ cost, for both either laminar and shear layer fluid conditions at the free stream velocity of 1.5 BL s^-1^. The concept of modelling EPOC for the locomotion gait was pioneered by Brett (*11*) in fish, which was applied to understand the thermal effects on locomotion costs of migratory salmon (*12*). The same non-aerobic cost framework is also used in sports science (*13*) (*14*). The sum of non-aerobic O_2_ cost and aerobic cost gives the total O_2_ cost. Based on the total O_2_ cost, we calculated total energy expenditure (TEE) by converting the total O_2_ cost to kJ • kg^-1^ using an oxy-calorific equivalent of 3.25 cal per 1 mg O_2_ (*15*).

### Analysis of force variation with angle of foil

Two-dimensional computational fluid dynamics (CFD) simulations were conducted to investigate the effects of foil angle of attack and the associated wake structures on the hydrodynamics of fish swimming in the wake. Experimental observations show that the fish body remains nearly stationary and inclined at a fixed angle relative to the incoming flow. Accordingly, an inclined, static NACA foil with the same size and cross-sectional shape as the fish body was employed to model a fish swimming downstream of a foil, as illustrated in the schematic in Fig. 8A. An in-house immersed-boundary-method (IBM)–based direct numerical simulation (DNS) solver was used to simulate fish swimming behind a foil at different angles of attack. A local mesh-refinement technique (*16*) was applied to accelerate the simulations. The Reynolds number was set to 5,000, based on the incoming flow velocity *U_∞_* and the foil chord length *c*. Two-layer of mesh refinement was used, with a finest grid size of Δ*_min_*= 3.25 × 10^,3^*c*. This resolution has been validated in previous studies (*17*) (*18*) to be sufficient for accurately resolving flow structures and achieving grid-independent results. Additional details of the numerical methods and simulation setup can be found in the previous study (*16*).

The simulations were performed over a duration of 60*T_c_*, where *T_c_* = *c*⁄*U_∞_* is the convective time scale based on the foil chord length, to ensure that the strongly nonlinear and highly unsteady flow at high angles of attack was fully developed and that statistically steady results were obtained. Time histories of drag coefficient (*C_D_*) and lift coefficient (*C_L_*) of fish swimming behind a foil fixed at different angles are shown in Figs. 8B and 8D. Live-fish experiments indicate that, at higher foil angles, the fish becomes more active in controlling its body posture and fins to maintain position and balance. This behaviour is hypothesized to be associated with intensified force fluctuations experienced by the fish when swimming within vortex wakes generated by the foil. To quantify these effects, we computed the root-mean-square (RMS) values of drag and lift coefficients to characterize their effective amplitude variations, and the RMS values of their time derivatives (*C_D_* and *C_L_*) to quantify how rapidly the force varies in time. The corresponding results, shown in Fig. S2, suggest that both drag and lift exhibit increasingly intense variations in both amplitude and time as the foil angle increases.

**Fig. S1.**
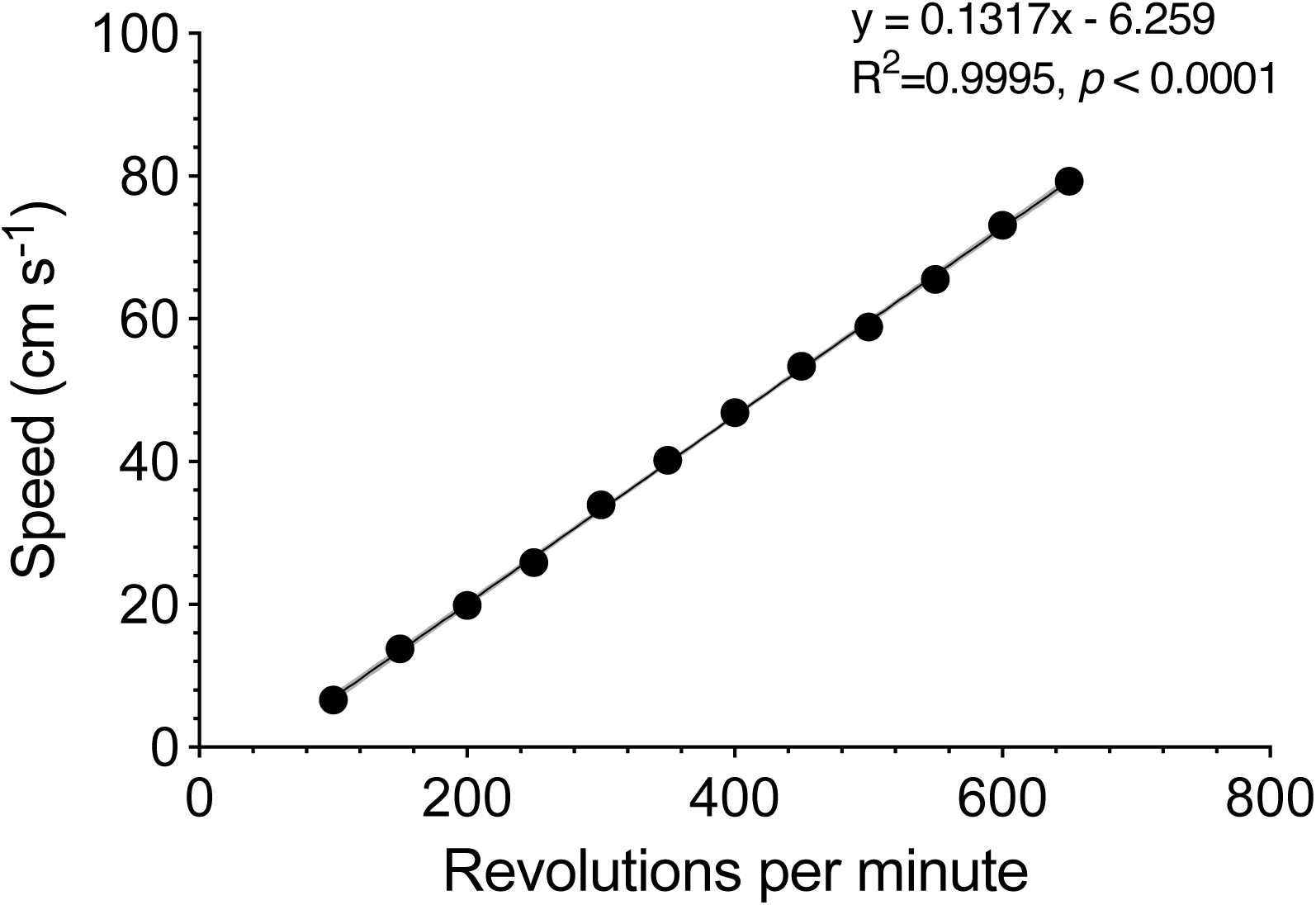
The mathematical regression between mean water velocity nd revolutions per minute of the propeller in the swim-tunnel respirometer. Particle image velocimetry (PIV) measured the flow field in the center of the swimming section of the swim-tunnel respirometer. The mean water velocity parallel to the flow direction is measured as speed on the y-axis. The revolutions per minute (RPM) is measured from the motor of the swim-tunnel respirometer. The motor drives a propeller inside the respirometer to move the water at different velocities. The measurement of twelve points enables the interpolation of the linear regression equation (R^2^ and *p* values) between speed and RPM of the motor (*see* figure legends).

**Fig. S2.**
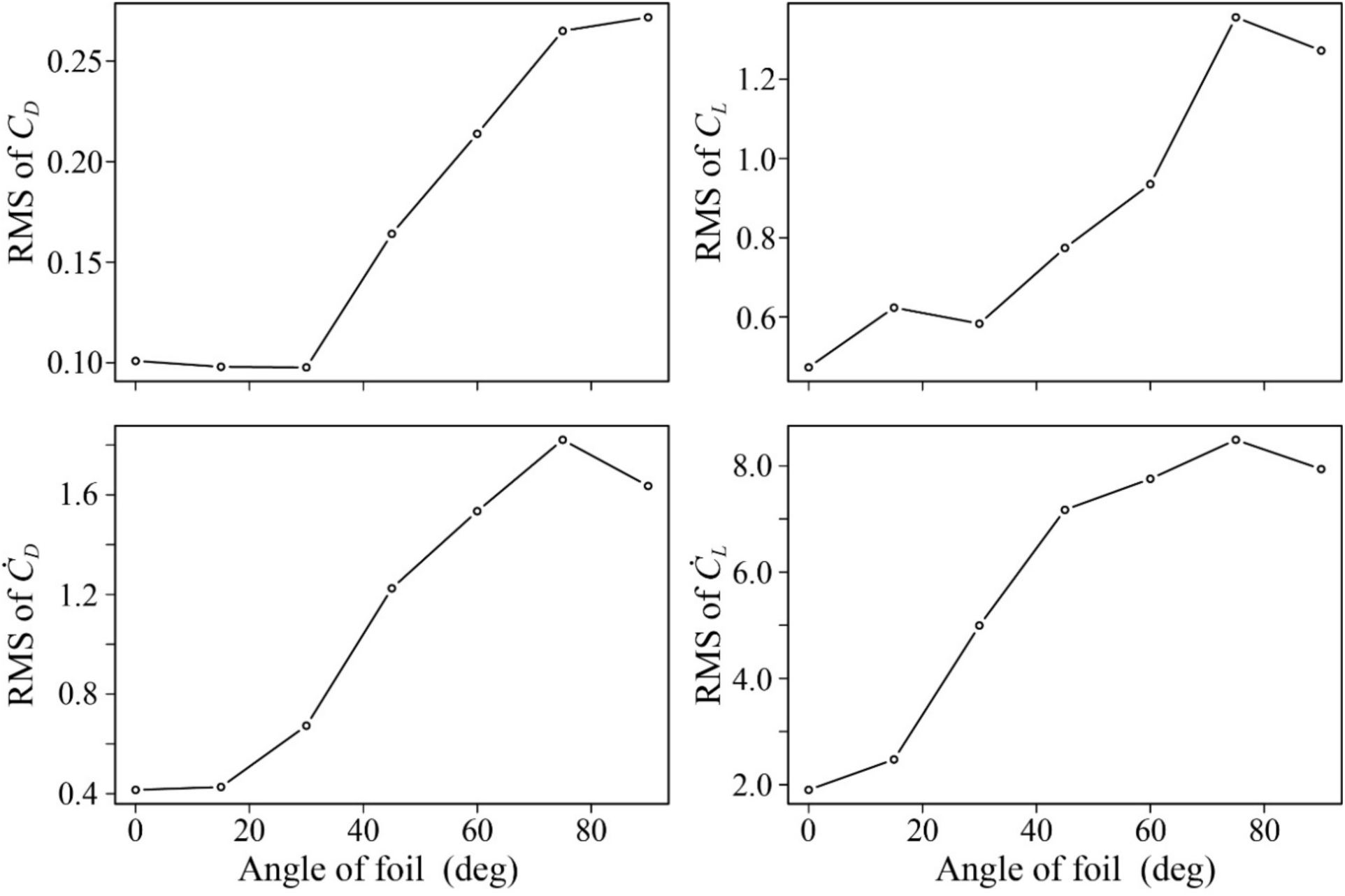
Variation in drag and lift computed from 2D CFD simulations experienced by a fish model swimming downstream of a foil fixed at different angles. The root-mean-square (RMS) values of the drag (A) and lift (B) coefficients are calculated to quantify the intensity of their amplitude variations. Larger RMS values indicate that the fish experiences larger effective variation in hydrodynamic forces. The RMS values of the time derivatives of the force coefficients, i.e., RMS values of ***C_D_*** (C) and ***C_L_*** (D), are computed to quantify the intensity of their temporal variations. Larger RMS values of the time derivatives indicate that the forces acting on the fish body vary more rapidly in time and are more strongly unsteady.

